# Serotonin Reduces Belief Stickiness

**DOI:** 10.1101/2023.12.08.570769

**Authors:** Vasco A. Conceição, Frederike H. Petzschner, David M. Cole, Katharina V. Wellstein, Daniel Müller, Sudhir Raman, Tiago V. Maia

**Author notes:** Address correspondence to:* Tiago V. Maia, Ph.D., Instituto de Medicina Molecular Faculdade de Medicina, Universidade de Lisboa, Av. Prof. Egas Moniz 1649-028 Lisboa, Portugal, Frederike H. Petzschner, Ph.D., Robert J. and Nancy D. Carney Institute for Brain Science Brown University, 164 Angell Street, Providence, RI, 02912, United States of America. Authors contributed equally (Vasco A. Conceição, Frederike H. Petzschner).

## Abstract

Serotonin fosters cognitive flexibility, but how, exactly, remains unclear. We show that serotonin reduces belief stickiness: the tendency to get “stuck” in a belief about the state of the world despite incoming contradicting evidence. Participants performed a task assessing belief stickiness in a randomized, double-blind, placebo-controlled study using a single dose of the selective serotonin reuptake inhibitor (SSRI) escitalopram. In the escitalopram group, higher escitalopram plasma levels reduced belief stickiness more, resulting in better inference about the state of the world. Moreover, participants with sufficiently high escitalopram plasma levels had less belief stickiness, and therefore better state inference, than participants on placebo. Exaggerated belief stickiness is exemplified by obsessions: “sticky” thoughts that persist despite contradicting evidence. Indeed, participants with more obsessions had greater belief stickiness, and therefore worse state inference. The opposite relations of escitalopram and obsessions with belief stickiness may explain the therapeutic effect of SSRIs in obsessive-compulsive disorder.

## INTRODUCTION

Serotonin fosters cognitive flexibility^1–3^, but the mechanisms through which it does so remain unclear. In this article, we propose and test a computational theory of a key mechanism through which serotonin fosters cognitive flexibility. The theory is that serotonin decreases the stickiness—i.e., the inflexibility or rigidity—of beliefs about the current state of the world. We demonstrate this effect empirically by combining a pharmacological manipulation in humans, a task designed to probe such stickiness, and a computational model that embodies our theoretical proposal.

Adequate inference about the current state of the world, and about when that state changes, is fundamental for flexible behavior. Humans, like other animals, adjust behavior to optimize the outcomes of their actions. In stable environments, behavior may often be optimized through the gradual learning of stimulus-response (S-R) associations^4,5^ via reinforcement learning^4,6,7^. Sometimes, however, contingencies change abruptly, and behavior must be adjusted quickly. One possible strategy in such cases is to increase the learning rate^8,9^. This solution, however, overwrites prior learning, which is suboptimal if the previous contingencies recur in the future. A more flexible solution is to infer that the world is now in a different (hidden) state, and learn new S-R associations for that state^10– 13^. Should the previous contingencies recur, one can infer that the world has returned to the previous state; all prior learning will then be available to guide behavior^13^.

A prototypical paradigm requiring state inference is reversal learning^13^. Reversal-learning experiments start with a set of contingencies that are later reversed. Many such experiments include multiple reversals, in which contingencies are changed back to prior configurations^1,14–17^. Participants who can properly infer the states, and state changes, in these tasks will perform better: as soon as they discover that a previous state (i.e., a previous configuration of contingencies) has been reinstated, they can adapt their behavior instantly.

Many studies^14,15,18,19^ (albeit not all^20^) suggest that performance following reversals depends on the orbitofrontal cortex (OFC), which is involved in state inference^11,13,21–25^. Such performance is also strongly modulated by serotonin^1,15,26–30^, particularly in the OFC^2,31–34^. Indeed, reduced serotonergic signaling in the OFC increases perseveration during reversal learning^1,2,31,32,35,36^, which led to the idea that prefrontal^1,26^, and especially orbitofrontal^2,33,34^, serotonin is important for cognitive flexibility. However, the mechanism through which orbitofrontal serotonin fosters flexibility in reversal learning—and more generally—is not fully understood.

We propose that orbitofrontal serotonin fosters cognitive flexibility by decreasing the stickiness of beliefs about the current state. This proposal derives from the intersection of two lines of evidence. First, the OFC represents beliefs about the current state^11,13,21– 25,37^. Second, biophysically detailed simulations show that serotonin decreases the tendency for neural representations in the OFC to become “sticky”—i.e., to form excessively strong attractors to which the network tends and from which it has difficulty escaping^33^. Combining these two lines of evidence, we arrive at our proposal that serotonin reduces the stickiness of beliefs about the current state (represented in OFC). A corollary of this proposal is that orbitofrontal serotonin is fundamental for flexible state inference—specifically, to detect state changes.

Reversal-learning experiments probe this ability to detect state changes, so our theory explains why orbitofrontal serotonin depletion causes perseveration in these experiments. When the contingencies change in these experiments, participants should infer that the state has changed. Under low serotonin, however, the state belief in OFC becomes “stuck” in the previous state, causing perseveration: responding according to the previous state.

Sticky orbitofrontal representations may also provide a natural explanation for obsessive-compulsive disorder (OCD)^33^. A second theoretical idea that we propose, and that we test preliminarily in this article, is that obsessions, in particular, result from sticky beliefs about state. The obsession “my hands are contaminated,” for example, can be seen as a sticky belief about the state of one’s hands that persists even after exhaustive handwashing. (As we explain in the Discussion, such belief need not be explicit and declarative; we use “belief” in a Bayesian sense.)

Several lines of evidence support the proposal that OCD may result from state stickiness^38,39^. First, as noted above, the OFC represents beliefs about the current state^11,13,21–25,37^; importantly, patients with OCD have prominent alterations in OFC structure and function^40–46^. Second, the conceptualization of obsessions as overly strong attractors to which network activity tends and from which it has difficulty escaping has considerable face validity^33,47,48^. Third, patients with OCD have atypical OFC activation during reversal learning^16,49–51^. Fourth, our proposal that obsessions result from sticky state representations in the OFC relates to our proposal that serotonin decreases such stickiness: the first-line pharmacological treatment for OCD consists of selective serotonin reuptake inhibitors (SSRIs)^52^, which increase extracellular serotonin^53^, normalize aberrant OFC activity in patients with OCD^40,54^, and may reduce the tendency of OFC networks to “get stuck”^33^.

In summary, we propose that serotonin and obsessions may relate to belief stickiness, and therefore to flexible state inference, in opposite directions: serotonin reduces belief stickiness, thereby facilitating flexible state inference; obsessions, conversely, are associated with increased belief stickiness, and therefore with inflexible state inference. To test these hypotheses, we developed a novel task and computational model to probe belief stickiness. We manipulated serotonin pharmacologically, administering a single dose of either the selective serotonin reuptake inhibitor (SSRI) escitalopram or a placebo to participants in a randomized, double-blind design (*Methods – Experimental protocol*). In addition, we measured participants’ escitalopram plasma levels when they performed the task (*Methods – Calculation of escitalopram plasma levels*). We assessed participants’ obsessionality using the Obsessive-Compulsive Inventory-Revised (OCI-R)^55^. Given that our hypotheses concerned specifically obsessions, we focused especially on the OCI-R’s “obsessions” subscale^55^.

We studied participants from the general population rather than patients with OCD because we wanted to investigate obsessionality as a dimensional construct that exists on a continuum^56,57^. We hypothesized that belief stickiness and flexible state inference would relate to obsessions (positively and negatively, respectively) along this continuum. Multiple lines of evidence support the usefulness of samples from the general population to investigate obsessive-compulsive symptoms dimensionally^58^.

## RESULTS

### Task

To probe belief stickiness and state inference, we developed a novel Go/NoGo reversal-learning task: the shell task (Fig. 1). Participants had to decide whether or not to collect different shells (Go or NoGo, respectively; Fig. 1a). Shells could be in one of three states (seasons): rewarding, punishing, or neutral (Fig. 1b). Each shell, however, alternated between only two of these states, with different shells alternating between different states (Fig. 1c). To adapt behavior quickly following state changes, especially when states recurred, participants had to constantly infer the state of each shell (Fig. 1d). Stickiness of beliefs about state caused difficulty adjusting behavior when states changed. The task had 216 trials (Fig. 1e).

**Fig. 1.**
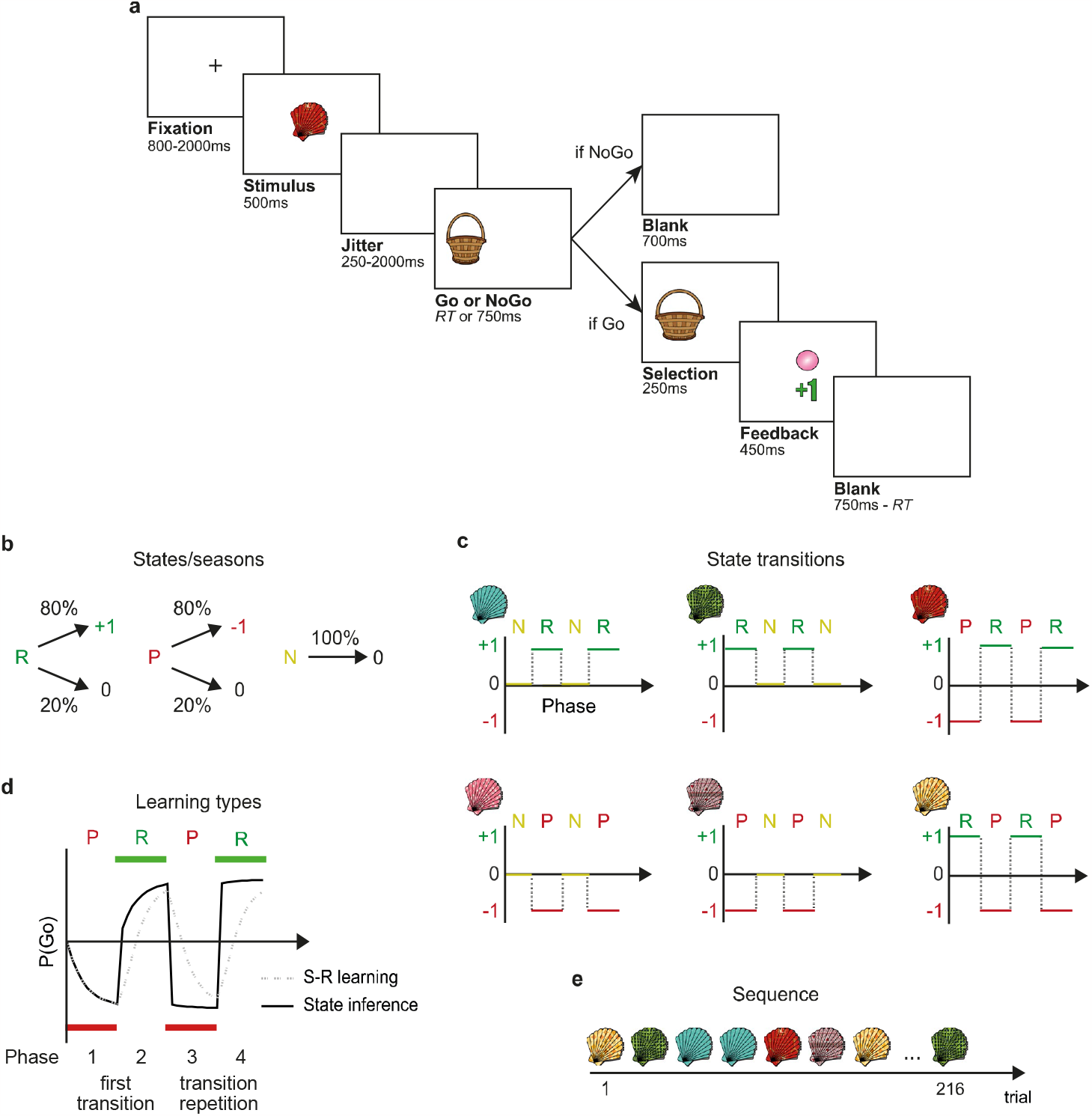
The novel Go/NoGo reversal-learning task: the shell task. **a**, Single trial: On each trial, participants were presented with one of six possible shells and had to decide whether to collect the shell to obtain its content (Go) or let it pass (NoGo). Shells were collected by pressing the button corresponding to the side of the presented bucket (Go); if the participant did not press that button within 750ms, the shell was not collected (NoGo). If the participant collected the shell, the bucket’s size increased momentarily, indicating that the shell was collected (*Selection* screen). Shells could contain a pearl (reward: +1 point), contain dirt (punishment: –1 point), or be empty (neutral: 0 points). If participants collected a shell, they were shown its content and obtained the corresponding points (*Feedback* screen); if they let a shell pass, they were not shown its content and did not gain or lose points. The duration of Go and NoGo trials was kept constant by adjusting the duration of the blank screen at the end of each Go trial. Participants were instructed to collect as many pearls as possible while avoiding collecting dirt (Methods Fig. 1); at the end of the task, they received a monetary reward proportional to their total points. **b**, Seasons (states): Across shells, there were three possible seasons, corresponding to three underlying (hidden) states. During a rewarding season (R), a shell would contain a pearl (+1 point) in 80% of the cases and be empty (0 points) in the remaining 20%. During a punishing season (P), a shell would contain dirt (–1 points) in 80% of the cases and be empty (0 points) in the remaining 20%. During a neutral season (N), the shell would always be empty (0 points). **c**, State transitions: Each shell alternated between two seasons (e.g., N and R, for the upper left shell), for a total of four phases, with a unique pattern of seasons per shell (e.g., NRNR for the same shell). There were six shells to encompass the three possible combinations of pairs of seasons (N and R, P and R, and N and P), with each of the three combinations being included in two shells, each of which started with a different season (e.g., NRNR and RNRN). Seasons varied independently for each shell. R and P seasons lasted 7-13 trials per shell; N seasons lasted 5-9 trials per shell. (N seasons were shorter because they were deterministic.) For each shell, the transition between the third and fourth phases was the same as the transition between the first and second phases (e.g., the transition from N to R for the NRNR shell). However, the transition between the third and fourth phases led to a previously visited season, whereas the transition between the first and second phases led to a new season. Participants were instructed that the probability of a given shell containing pearls, dirt, or nothing depended on its season, which could change throughout the task. No information was provided about the duration of seasons, their changes, or their reinforcement schedule, nor about potential differences between shells. Participants therefore had to learn which shells to collect at each time. **d**, Effect of state inference on task performance: The task was designed to characterize state inference, over and above simple stimulus-response (S-R) learning. This panel illustrates the differences in behavior between simple S-R learning (dashed gray) and the use of state inference (solid black), in which S-R learning occurs independently within each state. In the first phase, performance is largely indistinguishable in the two cases (note the overlapping curves for the first season) because state inference will tend to infer that the initial state remains the same, so all learning will be within that state. When a first state transition occurs—e.g., reversal from P to R in the figure—the participants’ responses might either change slowly, as new S-R learning overrides the old, or, if they are performing state inference and detect a change in state, change somewhat faster, as learning for the new state will occur unencumbered by the prior learning for the previous state (compare the dashed and solid lines during the first R season). The most notable divergence in behavior between simple S-R learning and the use of state inference, however, arises when a state recurs (compare the dashed and solid lines during phases three and four). In those situations, state inference supports better performance because it allows reutilization of the previously acquired knowledge about that state; simple S-R learning, in contrast, requires learning S-R associations all over again (and, moreover, starting from S-R associations that are no longer valid and hence need to be overridden). The best indicator for the use of state inference, without using a computational model, involves comparing performance when revisiting states that are preceded by identical state transitions—i.e., phases 2 and 4 (in this example, a comparison between the two R seasons, given that they are both preceded by the same state—the P season—but performance on the second R season will, with state inference, benefit from knowledge acquired in the first R season). Improved performance in phase 4 relative to phase 2 also distinguishes state inference from accounts that simply involve increasing learning rate(s) when states change because of the increased volatility^8^ or unexpected uncertainty^9^ during state transitions. Phases 2 and 4 are identical both in their contingencies and in how those contingencies differ from the contingencies in the preceding phase (phases 1 and 3, respectively). The changes in contingencies, and hence the volatility and unexpected uncertainty, are therefore similar in the transition to phase 2 and in the transition to phase 4; thus, these alternative accounts do not predict better performance in phase 4 relative to phase 2 (whereas the state-inference account does). **e**, Each participant completed 216 trials: 34 trials for each shell with N seasons (NRNR, NPNP, RNRN, and PNPN) and 40 trials for each of the remaining shells (PRPR and RPRP). Shells with N seasons were presented for fewer trials because, as noted above, N seasons were shorter because they were deterministic. The assignment of the six shell images to the six shell types (e.g., which shell image represented the NRNR shell and which image represented each of the other shell types), the exact duration and reinforcement schedule of each shell season, and the order of shell presentations differed pseudo-randomly across participants. Abbreviation: RT, reaction time.

### Participants

Participants were 50 men. The inclusion only of men is a limitation, but it allowed us to prevent confounds due to variation in serotonin metabolism^59^ and receptor availability^60^ during the menstrual cycle. We excluded 5 participants who did not understand the task, according to a debriefing questionnaire (*Methods – Debriefing questionnaire*), and 1 participant who selected Go in more than 90% of the trials. Of the remaining 44 participants (ages 24 ± 3 years), 20 and 24 were in the escitalopram and placebo groups, respectively. One of these participants did not fill out the OCI-R questionnaire and was therefore excluded from analyses that included that questionnaire.

### Participants performed state inference

We compared computational models from two families: one performed stimulus-response (S-R) learning; the other augmented S-R learning with a Bayesian state-inference layer (Fig. 2; Methods Fig. 2; *Methods – Computational models*). We call models in the latter family “S-S-R,” to highlight their hierarchical organization, with the state-inference layer atop the S-R layer (Fig. 2; Methods Fig. 2). S-R models learned to perform Go or NoGo for each shell. S-S-R models tried to infer each shell’s hidden state and learned to perform Go or NoGo within the shell’s inferred state.

**Fig. 2.**
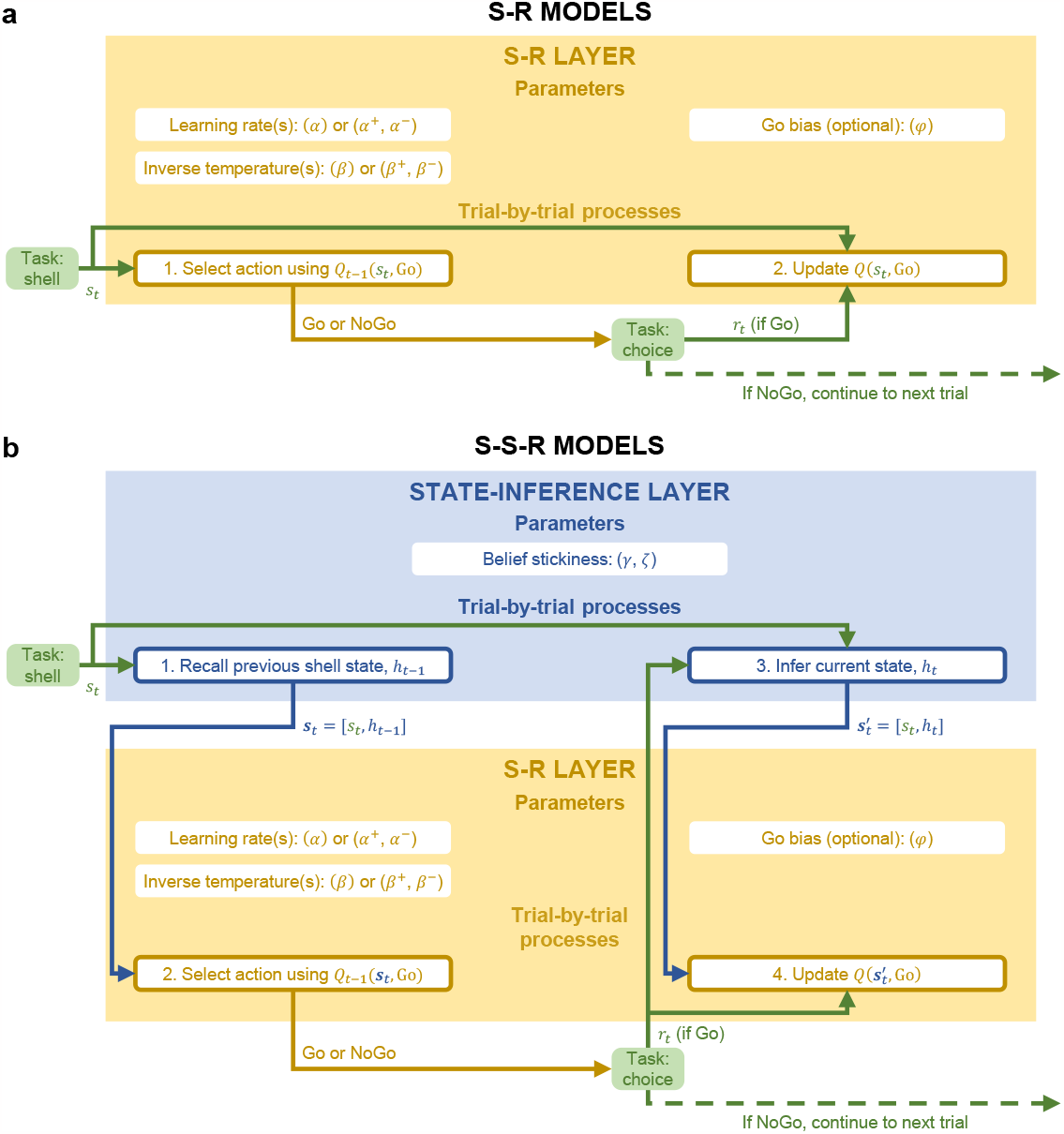
The model space was divided into two families: one implemented simple stimulus-response (S-R) learning (S-R models, panel a); the other augmented S-R learning with state inference (S-S-R models, panel b). **a**, S-R models only had an S-R layer. This layer implemented a variant of *Q*-learning^71^, with different models using different parameterizations to model aspects of S-R learning. Specifically, models differed in whether they (1) had one learning rate (*α*) or different learning rates for positive versus negative prediction errors (*α*^+^ and *α*^−^, respectively), (2) had one inverse temperature (*β*) or different inverse temperatures for positive and negative state-action weights (*β* ^+^ and *β* ^−^, respectively), and (3) did or did not include a Go bias (*φ*)^6,61–63^. *Q* values were learned only for Go; in our task, NoGo never resulted in feedback, so *Q*(*s*_*t*_,NoGo) was always 0. In S-R models, the state was univocally determined by the shell presented on the trial, *s*_*t*_. The S-R layer decided to perform a Go or NoGo by comparing *Q*(*s*_*t*_,Go) against the constant *Q*(*s*_*t*_,NoGo) of 0 (box 1). Then, if the action was a Go, the S-R layer updated *Q*(*s*_*t*_,Go) using the received reinforcement, *r*_*t*_ (box 2). **b**, S-S-R models added a state-inference layer atop the S-R layer. S-S-R models differed from S-R models only in the treatment of the state. Whereas in S-R models, the state was univocally determined by the shell presented on the trial, *s*_*t*_, S-S-R models tried to infer the shell’s hidden state (season), *h*. Thus, in S-S-R models, the state, ***s***_*t*_, consisted of an ordered pair: ***s***_*t*_= [*s*_*t*_, *h*]. At the beginning of the trial, when the shell was presented, the state-inference layer considered the previously inferred state of the shell, *h*_*t*–1_, as the best bet for the shell’s current state (box 1). (The subscript “*t*” denotes the *t*^th^ presentation of the shell, not the overall trial number; thus, the subscript “*t*–1” denotes the previous presentation of the shell.) The state-inference layer provided the pair ***s***_*t*_= [*s*_*t*_, *h*_*t*–1_] to the S-R layer, which decided to perform a Go or NoGo using *Q*(***s***_*t*_,Go) (box 2). If the action selected was Go, this resulted in a reinforcement, *r*_*t*_. The state-inference layer then used this observed reinforcement to try to infer the shell’s current state, *h*_*t*_ (box 3), which might or not differ from *h*_*t*–1_. This process was modulated by the belief-stickiness parameters, *γ* and *ζ*. Briefly, larger values of *γ* and *ζ* led to “stickier” beliefs—i.e., a greater tendency to perseverate on the prior beliefs, thereby hampering the ability to detect state changes (see *Methods* – *Computational models* – *S-S-R models* and *Supplementary results – Effects of γ and ζ on task performance*). Having inferred *h*_*t*_, the state-inference layer then provided the pair 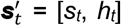 to the S-R layer. The S-R layer then updated the *Q*-value for the shell’s newly inferred state, 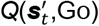, using the received reinforcement, *r*_*t*_ (box 4). This ensured that learning occurred for the newly inferred state. The role of the state-inference layer was to provide top-down information about state to the S-R layer, for use in action selection (***s***_*t*_) and state-action learning 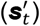. Through this arrangement, S-S-R models had independent *Q*-values not only for different shells (like S-R models did) but also for different hidden states for each shell. Methods Fig. 2 shows the processes depicted here in more detail (including the steps involved in state inference and all model equations).

We implemented eight partially nested S-R models, with up to five parameters characterizing variations of S-R learning (*Methods* – *Computational models* – *S-R models*)^6,61–63^. Similarly, we implemented eight S-S-R models, with each S-S-R model augmenting the corresponding S-R model with the state-inference layer. The S-S-R models share some similarities with prior models^64,65^ but were developed specifically for this study. S-S-R models had two additional parameters, *γ* and *ζ*, designed to tap into the underlying construct of belief stickiness through complementary mechanisms (*Methods – Computational models* – *S-S-R models*). Larger values of each of *γ* and *ζ* corresponded to greater belief stickiness (i.e., to a greater tendency to stick with the previously inferred state even when incoming evidence suggested a state change).

We fit models using thermodynamic integration (*Methods – Model inversion*), which adequately penalizes model complexity^66,67^. We could accurately recover models and parameters (*Supplementary results – Model- and parameter-recovery simulations*). When we generated simulated data using simple models, we accurately recovered those simple models, showing that our model-selection pipeline adequately penalized model complexity (*Supplementary results – Model- and parameter-recovery simulations*).

The most complex S-S-R model, S-S-R-8 (*Methods – Computational models*), was the winning model (Fig. 3a–b; protected exceedance probability^68^ ∼ 1), showing that participants performed state inference (as has been found in prior reversal-learning tasks^69,70^). The model frequencies did not differ between the escitalopram and placebo groups (posterior probability of equal frequencies = 0.986). Model S-S-R-8 fit participants’ individual trial-by-trial choices well [average (±SD) McFadden’s pseudo-*R*^2^: 0.35 (±0.14); see Fig. 4 for an example fit].

**Fig. 3.**
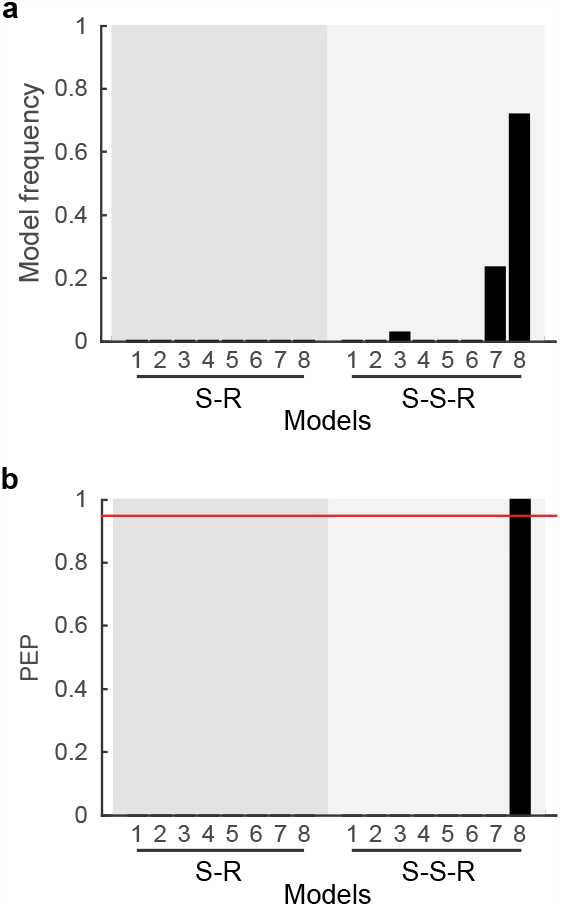
Behavior was best explained by a model that extends stimulus-response learning with state inference. **a**, Frequencies of the different candidate models, calculated as the posterior means of the model frequencies in the population, estimated with Bayesian model comparison. Models are grouped into simple stimulus-response (S-R) models and models that extend S-R models with a higher-level state-inference layer (S-S-R models). For a description of the various models, see *Methods – Computational models*. Model S-S-R-8 had the highest frequency. **b**, Protected exceedance probabilities (PEPs) for the different S-R and S-S-R models. The PEP for each model corresponds to the probability that that model is more frequent than any other model, correcting for the possibility that that finding was due to chance^68^. The red line indicates the threshold for confident selection of a model: a PEP above 0.95^68^. Model S-S-R-8 was confidently selected as the best model.

**Fig. 4.**
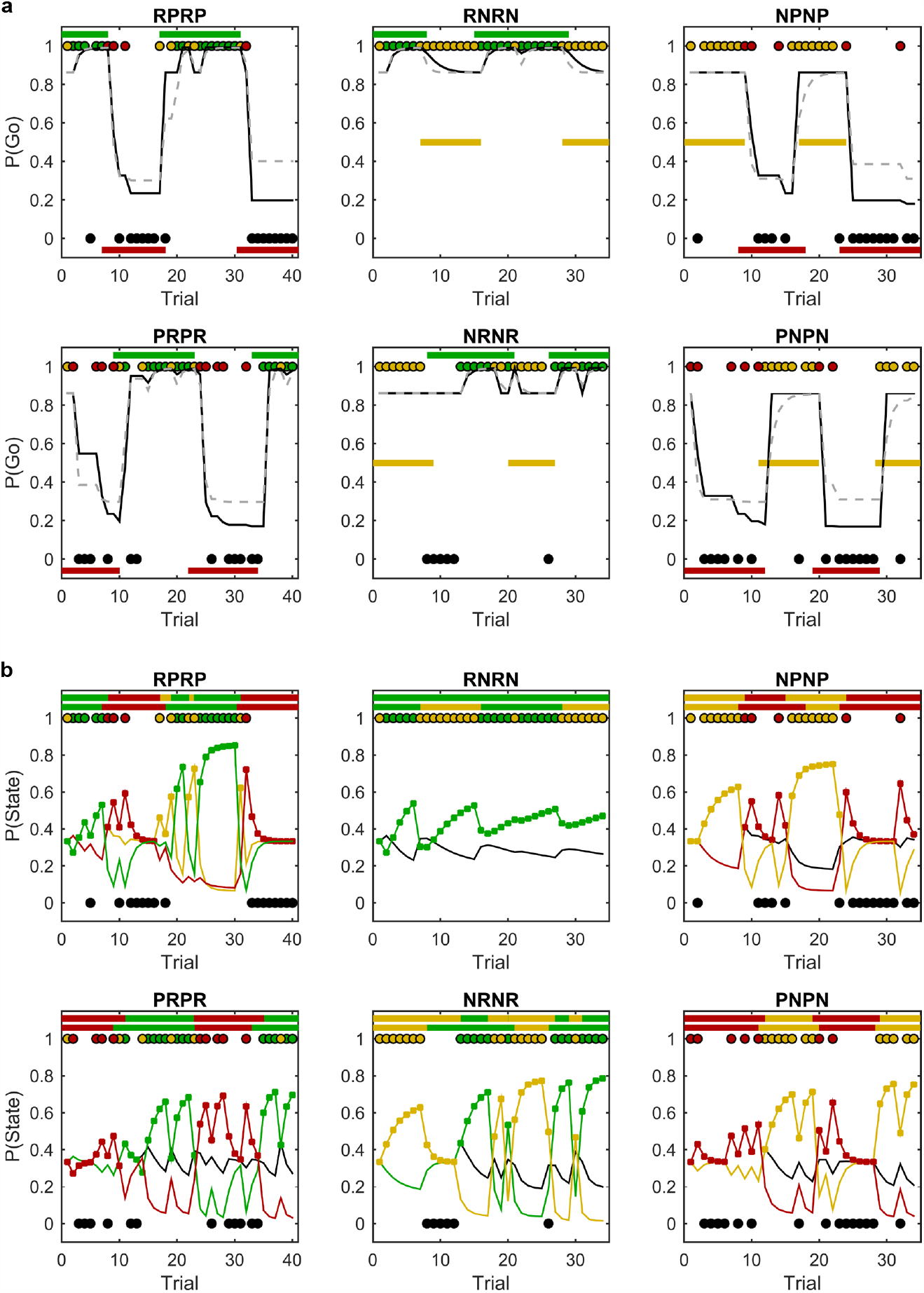
Model fit and behavioral data for an example participant. Each plot represents a shell. Points at the bottom and top of each plot indicate the behavioral response of the participant on each trial: black points at the bottom represent NoGo responses; colored points at the top represent Go responses, with the specific color depicting the reinforcement on the trial (green: reward; yellow: neutral; red: punishment). **a**, Probability of performing a Go response, P(Go), based on the fit of the winning model (S-S-R-8; solid black line), which includes state inference, and its corresponding S-R model (S-R-8; dashed gray line), which does not. Horizontal color-coded bars indicate the underlying season of the shell (green: rewarding; yellow: neutral; red: punishing). The winning model is more in line with the participant’s behavior, particularly when seasons recur (last two seasons for each shell), when previously learned information about those seasons can be reused. **b**, Belief trajectory of the winning model, S-S-R-8, for the participant. Horizontal color-coded bars on top indicate the state inferred by the model (with rewarding, neutral, and punishing inferred states depicted in green, yellow, and red, respectively). The true underlying states are plotted immediately below, to facilitate comparison between the true and inferred states. The plotted lines depict the posterior probability distribution of the beliefs about the shell state, P(state), on each trial: the beliefs that the shell is in rewarding, neutral, and punishing states are represented in green, yellow, and red, respectively. Note that, in the model, each state corresponds to a distribution of outcomes; states are not labeled *a priori* as representing rewarding, neutral, or punishing seasons. To assign such labels to states—to represent the inferred states in the top horizontal bars and the beliefs in the plotted lines—we assigned rewarding, neutral, and punishing labels to states whose final outcome distribution consisted mostly of reward, neutral, or punishment outcomes, respectively. Given that the task included three season types, for simplicity, the model assumed that there could be up to three hidden states (*Methods* – *S-S-R models*). In all plots except the one at the top left, the model only inferred up to two states; it therefore never associated any outcomes with the third state, which we therefore represent in black (as it does not have a predominance of any outcome type). Note also that some lines overlap and therefore are not always visible: this occurs in the top middle panel, in which the model only inferred one state, and there are therefore two states for which nothing was ever learned, so there are two overlapping black lines; it also occurs, for example, in the initial trials in each plot because, as long as the model has only inferred one state, the beliefs for the other two states are equal. The markers on the plotted lines indicate the state that the model believes applies after each trial; usually, this corresponds to the state with the largest posterior probability, but in some cases it does not because the model has a (parameter-dependent) tendency to continue to stick with the prior belief (*Methods – S-S-R models*).

Model S-S-R-8 also replicated participants’ group-level behavior (Fig. 5). Participants learned the seasons—the probability of performing a Go response was largest in rewarding seasons, intermediate in neutral seasons, and smallest in punishing seasons— and the model captured this effect (Fig. 5a). In addition, participants’ choices changed as seasons changed, and the model also captured these changes (Fig. 5b). These two effects, however, can also be captured using only S-R learning: model S-R-8, the S-R equivalent of S-S-R-8, captured them (Extended Data Fig. 1a–b). To assess specifically whether participants and model S-S-R-8 adequately inferred the hidden states (seasons), we compared performance when a state recurred, following the same state transition— i.e., we compared phases 2 and 4 of each shell. Adequate state inference would be reflected in better performance in phase 4, when the state recurred, than in phase 2, when the state first occurred (Fig. 1d). Participants indeed performed better in phase 4 than in phase 2, and model S-S-R-8 captured this effect (Fig. 5c). Model S-R-8, which did not include state inference, could not capture this effect (Extended Data Fig. 1c).

**Fig. 5.**
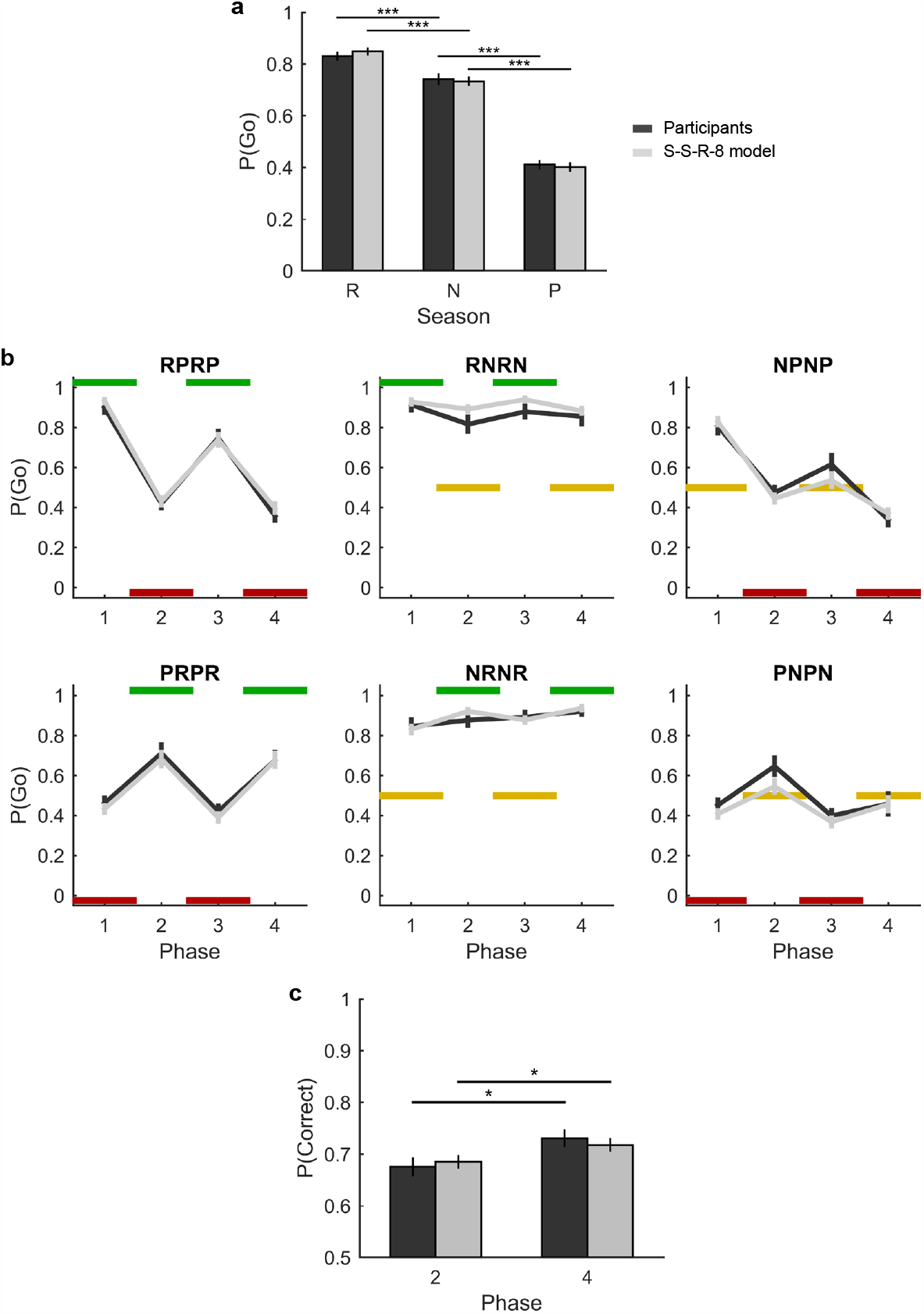
The winning model, S-S-R-8, replicates participants’ group-level behavior. The average of participants’ behavior is shown in dark gray; the average of the model fits for individual participants is shown in light gray. **a**, Average (±SEM) of the probability of Go responses, P(Go), exhibited by participants (left bar of each pair; dark gray) and model fits (right bar of each pair; light gray) for each season type: rewarding (R), neutral (N), and punishing (P). We analyzed both participants’ behavior and the model fits using a generalized linear mixed-effects model with a logit link, with P(Go) as the dependent variable, and with season type (rewarding, neutral, or punishing) as the independent variable (and with the intercept as a random effect to capture individual propensities to do Go). P(Go) was largest for rewarding, intermediate for neutral, and smallest for punishing seasons, for both participants (main effect of season type: 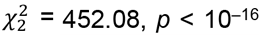; repeated contrasts: rewarding vs. neutral, *b* = 0.56, *z* = 5.05, *p* < 10^−6^, 95% CI [0.34, 0.78], *OR* = 1.77; neutral vs. punishing, *b* = 1.45, *z* = 16.55, *p* < 10^−16^, 95% CI [1.28, 1.62], *OR* = 4.25) and model fits (main effect of season type: 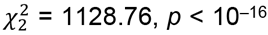; repeated contrasts: rewarding vs. neutral, *b* = 0.71, *z* = 9.32, *p* < 10^−16^, 95% CI [0.56, 0.86], *OR* = 2.03; neutral vs. punishing, *b* = 1.47, *z* = 25.51, *p* < 10^−16^, 95% CI [1.35, 1.58], *OR* = 4.34). **b**, Average (±SEM) P(Go) as a function of the shell phases for each of the six shells, for participants (dark gray) and model fits (light gray). Both participants and model fits tracked the seasons, with P(Go) increasing in R seasons (green) and decreasing in P seasons (red) for all shells. Behavior in N seasons (yellow) was more variable, but predictably so: both participants and the model fits tended to have substantially larger P(Go) in N seasons for shells in which the N seasons alternated with R seasons (RNRN and NRNR), as collecting shells in those cases never had any adverse consequences, than for shells in which the N seasons alternated with P seasons (PNPN and NPNP), as in those cases collecting the shell in the wrong season could lead to a loss. The model fits tracked participants’ behavior closely. The only difference between the behavior of the model and that of participants is that the model seems to have had greater difficulty identifying transitions to N seasons the first time that these occurred. This difficulty can be seen in phase 2 in the upper middle plot, phase 3 in the upper right plot, and phase 2 in the lower right plot; it also explains the better performance of the model relative to participants in phase 3 in the upper middle plot (because, by having difficulty identifying the N season in phase 2, the model was more likely to still be in season R at the onset of phase 3). Transitions to N seasons are more difficult to detect because R and P seasons also include neutral outcomes. **c**, Average (± SEM) probability of a correct response, P(Correct), in the phases following the first and second identical shell-state transitions—phases 2 and 4, respectively—for participants (dark gray) and model fits (light gray). P(Correct) was calculated using only R and P seasons, as there was no correct response for N seasons. P(Correct) was greater in phase 4 than in phase 2, for both participants (mean difference = 0.05, *t*_43_ = 2.61, *p* = .012, 95% CI [0.01, 0.10], Cohen’s *d* = 0.39) and model fits (mean difference = 0.03, *t*_43_ = 2.60, *p* = .013, 95% CI [0.01, 0.06], Cohen’s *d* = 0.39). \**p* < .05. \*\*\**p* < .001.

We verified that the conclusion that participants performed state inference was robust by implementing two additional model families to exclude alternative explanations (*Methods – Computational models – Alternative model families*). First, a possible concern with our task is that if participants get better at the task over time—i.e., if they start to learn faster— even without performing state inference, performance will increase later in the task, when states are revisited, which could mask as state inference. We therefore implemented a model family without state inference but with learning rates that could vary across the task. Second, several studies have related serotonin, OCD, and obsessive-compulsive symptoms to “stimulus stickiness”: the tendency to continue to choose the same stimulus^72–75^. Stimulus stickiness is confounded by state inference (see Discussion); to disentangle these processes, we implemented a model family without state inference but with stimulus stickiness. Model S-S-R-8 was still confidently selected when considering these two additional families (protected exceedance probability = 0.999; Extended Data Fig. 2).

A third computational alternative is that participants simply increased the learning rate(s) when states changed due to increased volatility^8^ or unexpected uncertainty^9^. That alternative, however, would have difficulty capturing the increase in performance from phase 2 to phase 4 (Fig. 5c) because the changes in contingencies, and therefore the volatility and unexpected uncertainty, are similar in both cases (Fig. 1d).

### The two state-inference parameters tapped into the same underlying construct of belief stickiness

Consistent with our hypothesis that *γ* and *ζ* tapped into a single underlying belief-stickiness construct, these parameters correlated strongly (*r*_42_ = .59, *p* < 10^−4^). Our primary interest was how this belief-stickiness construct relates to escitalopram (plasma level and group) and OCI-R scores (particularly, obsessionality). To analyze this relation, we used canonical correlation analysis (CCA; *Methods – Statistical analyses – Canonical correlation analysis*). We set *γ* and *ζ* as the *y* variables; we set escitalopram plasma level, group, OCI-R obsessing, and OCI-R other as the *x* variables (where we defined “OCI-R other” as the sum of all OCI-R subscales other than the OCI-R obsessing subscale). We split OCI-R scores into OCI-R obsessing and OCI-R other because, as noted in the Introduction, our hypotheses concerned specifically obsessions, so we were especially interested in the relations to OCI-R obsessing (but we did not want to exclude the other subscales from the analysis). The OCI-R obsessing subscale also best distinguishes patients with OCD from non-anxious controls^55^.

The CCA was significant (Wilks’s Λ = 0.66, *F*_8,74_ = 2.18, *p* = .039). The canonical loadings for *γ* and *ζ* on the first canonical variate for the *y* variables were both positive (0.999 and 0.636, respectively); this first canonical variate therefore captured our hypothesized latent belief-stickiness construct. The first canonical correlation was 0.504. The second pair of canonical variates was not significant (root 2: Wilks’s Λ = 0.88, *F*_3,38_ = 1.76, *p* = .171), suggesting that the latent belief-stickiness construct captured all structure in *γ* and *ζ* relevant for their relation to escitalopram (levels or group) and OCI-R scores (obsessing and other).

### Escitalopram plasma levels and obsessions related negatively and positively, respectively, to belief stickiness

As noted in the Introduction, our theoretical proposals are that serotonin reduces belief stickiness and that obsessions are associated with increased belief stickiness. We therefore hypothesized that escitalopram plasma levels and OCI-R obsessing scores related negatively and positively, respectively, to belief stickiness. Consistent with these hypotheses, the cross-loadings of escitalopram plasma level and OCI-R obsessing scores on the first canonical variate for *γ* and *ζ* (which capture the relations of escitalopram level and OCI-R obsessing, respectively, with the latent belief-stickiness construct) were negative (–0.355, *p* = .019, Fig. 6a) and positive (0.359, *p* = .017, Fig. 6b), respectively. The cross-loading of OCI-R other on that first canonical variate was also positive (0.215) but not significant (*p* = .165).

**Fig. 6.**
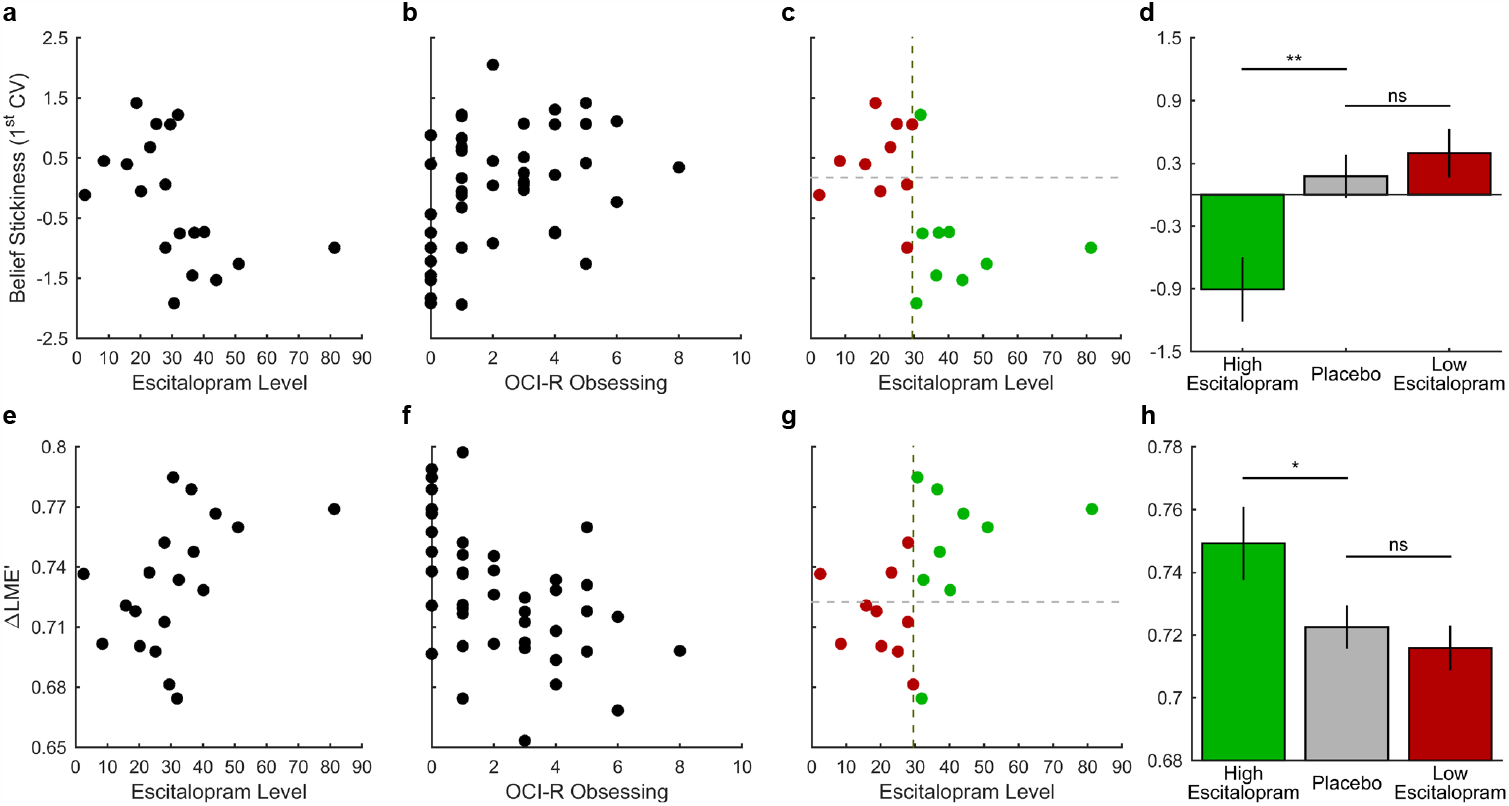
Higher escitalopram levels and higher obsessive traits had opposite effects, both on belief stickiness (top row) and on state inference (bottom row). Obsessive traits, belief stickiness, and state inference were assessed, respectively, by OCI-R obsessing scores, scores on the first canonical variate (1^st^ CV) for γ and *ζ*, and ΔLME’ (a Box-Cox transformation of the difference between the log model evidence for the selected model, S-S-R-8, and that for its corresponding S-R model, S-R-8; see text). Higher escitalopram levels decreased belief stickiness (panel a), thereby improving state inference (panel e); higher obsessive traits had the opposite effect, increasing belief stickiness (panel b) and thereby hindering state inference (panel f). A median-split of the escitalopram group into participants with high and low levels of escitalopram shows that only sufficiently high levels of escitalopram reduced belief stickiness relative to placebo (panels c-d), so only sufficiently high levels of escitalopram improved state inference relative to placebo (panels g-h). **a**, Higher escitalopram levels decreased belief stickiness. **b**, Higher obsessive traits increased belief stickiness. **c**, Only sufficiently high escitalopram levels reduced belief stickiness. This panel presents the same data as panel (a), but it also shows the mean belief stickiness for the placebo group (horizontal dashed line) and a median split on escitalopram levels (dashed vertical line). To aid in visualization, participants below and above the median split are color-coded in red and green, respectively. All participants above the median split, except one, had belief stickiness smaller than the mean belief stickiness for the placebo group (i.e., all participants color-coded green, except one, were placed in the lower-right quadrant). Most participants below the median split had belief stickiness greater than the mean belief stickiness for the placebo group (i.e., most participants color-coded red were in the upper-left quadrant). **d**, Belief stickiness was not the same across high-escitalopram, placebo, and low-escitalopram groups (one-way ANOVA: *F*_2,40_ = 6.02, *p* = .005, partial *η*^2^ = 0.23). Belief stickiness was smaller in the high-escitalopram group than in the placebo group (mean difference = –1.08, *t*_40_ = –3.08, *p* = .004, 95% CI [–1.79, –0.37], Cohen’s *d* = –1.20). The placebo and low-escitalopram groups, in contrast, did not differ significantly (mean difference = –0.22, *t*_40_ = –0.66, *p* = .515, 95% CI [–0.91, 0.46], Cohen’s *d* = –0.25). Shown are mean (±SEM) belief-stickiness values. The high- and low-escitalopram groups consist of participants in the escitalopram group who were above and below the median split on escitalopram levels, respectively. **e**, Higher escitalopram levels improved state inference. **f**, Higher obsessive traits hindered state inference. **g**, Only sufficiently high escitalopram levels improved state inference. This plot relates to panel (e) in the same way that panel (c) relates to panel (a). It shows the mean ΔLME’ for the placebo group (horizontal dashed line) and a median split on escitalopram levels (dashed vertical line). To aid in visualization, participants below and above the median split are color-coded in red and green, respectively. All participants above the median split, except one, had ΔLME’ values larger than the mean ΔLME’ for the placebo group (i.e., all participants color-coded green, except one, were placed in the upper-right quadrant), indicating better state inference. Most participants below the median split had ΔLME’ values smaller than the mean ΔLME’ for the placebo group (i.e., most participants color-coded red were in the lower-left quadrant), indicating worse state inference. **h**, ΔLME’ was not the same across high-escitalopram, placebo, and low-escitalopram groups (one-way ANOVA: *F*_2,40_ = 3.27, *p* = .048, *η*^2^ = 0.14). ΔLME’ was larger in the high-escitalopram group than in the placebo group (mean difference = 0.027, *t*_40_ = 2.23, *p* = .032, 95% CI [0.002, 0.051], Cohen’s *d* = 0.87). The placebo and low-escitalopram groups, in contrast, did not differ significantly (mean difference = 0.007, *t*_40_ = 0.58, *p* = .566, 95% CI [–0.017, 0.030], Cohen’s *d* = 0.22). Shown are mean (±SEM) ΔLME’ values. The high-and low-escitalopram groups consist of participants in the escitalopram group who were above and below the median split on escitalopram levels, respectively. \**p* < .05. \*\**p* < .01. ns: not statistically significant.

### Only sufficiently high escitalopram levels decreased belief stickiness relative to placebo

The cross-loading of group on the first canonical variate for *γ* and *ζ* represents a point biserial correlation that tests whether groups differed on the latent belief-stickiness construct. Consistent with the idea that the escitalopram group might have lower belief stickiness than the placebo group, this cross-loading was negative (–0.199); however, it was not significant (*p* = .200).

The absence of a significant group effect seems surprising considering the significant effect of escitalopram level. These seemingly contradictory findings can be reconciled, however, if only sufficiently high levels of escitalopram decrease belief stickiness relative to placebo. Low escitalopram levels might have no effect or could even increase belief stickiness because whereas high acute SSRI doses tend to increase cortical serotonin, low acute SSRI doses do not increase cortical serotonin and may even decrease it (through mechanisms that we discuss in the Discussion)^76^.

Consistent with this idea, a median split of the escitalopram group into low-vs. high-escitalopram levels showed that, in the high-escitalopram group, all participants except one had belief stickiness (i.e., scores on the first canonical variate for *γ* and *ζ*) below the mean belief stickiness for the placebo group (Fig. 6c). We therefore compared belief stickiness across the high-escitalopram, placebo, and low-escitalopram groups (after confirming that model frequencies did not differ between these groups: posterior probability of equal frequencies ∼ 1). As we had hypothesized, belief stickiness was smaller in the high-escitalopram group than in the placebo group, whereas the placebo and low-escitalopram groups did not differ significantly (Fig. 6d). Thus, escitalopram decreases belief stickiness relative to placebo, but only if the escitalopram level is sufficiently high.

### Escitalopram plasma levels and obsessions related positively and negatively, respectively, to state inference

As noted in the Introduction, our theoretical proposals are that serotonin reduces belief stickiness, thereby facilitating state inference, and that obsessions are associated with increased belief stickiness, which hinders state inference. We showed above that escitalopram plasma level and OCI-R obsessing scores related to belief stickiness as predicted by these proposals: negatively and positively, respectively. Next, we tested if escitalopram plasma level and OCI-R obsessing scores also related to state inference as predicted by these proposals: positively and negatively, respectively.

We quantified state inference for each participant by calculating the difference between the log model evidence (LME) for the selected model, S-S-R-8, and that for its corresponding S-R model (S-R-8). This difference, ΔLME, indicates the extent to which the state-inference layer, present in model S-S-R-8 but not in model S-R-8, is necessary to explain the participant’s behavior, so it captures the extent to which the participant’s behavior was based on (adequate) state inference. To assess how ΔLME depended on escitalopram (level or group) and OCI-R scores, we regressed ΔLME on group, escitalopram level, OCI-R obsessing, and OCI-R other (see *Methods – Statistical analyses – Coding of group and escitalopram plasma level in regressions* for details on how group and escitalopram level were coded).

The residuals of this regression were heteroscedastic (Breusch-Pagan test using the fitted values: 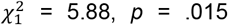), so we transformed ΔLME using a Box-Cox transformation. With the transformed value, ΔLME’ (*λ*_1_ = –1.20, *λ*_2_ = 4.61), there was no longer evidence for heteroscedasticity of the residuals (Breusch-Pagan test using the fitted values: 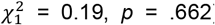), so we conducted the regression using ΔLME’. Escitalopram level and OCI-R obsessing related positively and negatively, respectively, to ΔLME’ (*b* = 0.0008, *t*_38_ = 2.19, *p* = .035, 95% CI [0.0001, 0.0016], Fig. 6e, and *b* = – 0.0093, *t*_38_ = –3.26, *p* = .002, 95% CI [–0.0151, –0.0035], Fig. 6f, respectively), showing that, as expected, higher escitalopram levels and higher obsessive traits facilitated and hindered state inference, respectively (presumably because, as shown above, they decrease and increase belief stickiness, respectively). Consistent with all other analyses in the article, neither group nor OCI-R other related significantly to ΔLME’ (*b* = 0.0045, *t*_38_ = 0.53, *p* = .600, 95% CI [–0.0127, 0.0217], and *b* = 0.0008, *t*_38_ = 0.97, *p* = .336, 95% CI [–0.0009, 0.0025], respectively).

We replicated the positive relation between escitalopram plasma level and state inference using simpler (albeit cruder) analyses that did not rely on computational models (*Supplementary results – Replication of the positive association between escitalopram plasma level and state inference*).

### Only sufficiently high escitalopram levels facilitated state inference relative to placebo

In the belief-stickiness analysis, group did not have a significant effect on belief stickiness, and follow-up analyses showed that this lack of significance occurred because only sufficiently high escitalopram levels—defined as escitalopram levels above the median— decreased belief stickiness relative to placebo (Fig. 6c-d). We hypothesized that the lack of a significant group effect on ΔLME’ in the previous section occurred, similarly, because only sufficiently high escitalopram levels increased ΔLME’. Consistent with this hypothesis, the median split of the escitalopram group into low-vs. high-escitalopram levels showed that, in the high-escitalopram group, all participants except one had ΔLME’ values above the mean ΔLME’ for the placebo group (Fig. 6g). We therefore compared ΔLME’ across the high-escitalopram, placebo, and low-escitalopram groups. Consistent with our hypothesis and with the equivalent analysis of belief stickiness, ΔLME’ was larger in the high-escitalopram group than in the placebo group, whereas the placebo and low-escitalopram groups did not differ significantly. Taken together, the findings with a median split of the escitalopram group showed that only sufficiently high escitalopram levels decreased belief stickiness (Fig. 6c-d), thereby facilitating state inference (Fig. 6g-h), relative to placebo.

### Neither escitalopram nor obsessive traits related to S-R parameters

We also tested if non-state-related (i.e., S-R) parameters related to escitalopram (level or group) and/or OCI-R obsessing scores. For that purpose, we used a multivariate regression, with the S-R parameters—*α*^+^, *α*^−^, *β*^+^, *β*^−^, and *φ*—as the vector of dependent variables, and with group, escitalopram level, OCI-R obsessing scores, and OCI-R other as the independent variables. This regression was not significant (Wilks’s Λ = 0.59, *F*_20,113.72_ = 0.98, *p* = .486), so there was no evidence that the S-R parameters related to escitalopram (level or group) or OCI-R scores. Moreover, the regression’s adjusted multivariate *R*^2^, calculated using a formula that generalizes the adjusted *R*^2^ to the multivariate case^77^, was negative (adjusted multivariate *R*^2^ = –0.01).

If indeed escitalopram (level and group) has no effect on the S-R parameters, that has two corollaries. First, it suggests that escitalopram did not relate to the Go bias (which was one of the S-R parameters, *φ*). This result would go against an earlier report that citalopram, which is a mixture of 50% S-citalopram (escitalopram) and 50% R-citalopram, increased the Go bias in a simpler task^78^. We therefore conducted an additional analysis investigating whether escitalopram related to the Go bias. That analysis confirmed that escitalopram predominantly, if not selectively, affected state inference rather than the Go bias (*Supplementary results – Escitalopram did not relate to S-R parameters – Escitalopram plasma levels predominantly affected state inference rather than a Go bias*). Second, it suggests that escitalopram did not relate to punishment-based learning or performance, which should be captured in α^−^ or β^−^, respectively. Given the longstanding ideas relating serotonin to punishment^79–81^, we conducted additional analyses investigating whether escitalopram related to punishment. Those analyses also found no evidence that escitalopram modulated punishment-based learning or performance (*Supplementary results – Escitalopram did not relate to S-R parameters*).

Of course, these arguments are based on arguing for the null hypothesis, which is fraught with difficulties. Still, the nearly 0 and even negative adjusted *R*^2^ in the multivariate regression of the S-R parameters on escitalopram (level and group) and OCI-R scores suggests that, if escitalopram (level or group) and/or OCI-R scores (obsessive or other) relate to S-R parameters at all, such relations are likely weak. The effect sizes in the aforementioned supplemental analyses lead to a similar conclusion (*Supplementary results – Escitalopram did not relate to S-R parameters*). Overall, then, escitalopram levels and OCI-R obsessing scores selectively, or at least preferentially, affected belief stickiness.

## DISCUSSION

We found that higher escitalopram plasma levels were associated with lower stickiness of beliefs about state and better state inference. Moreover, participants with sufficiently high escitalopram plasma levels—specifically, participants with escitalopram plasma levels above the median—had lower belief stickiness and better state inference than participants on placebo. These findings support our first theoretical proposal, which is that serotonin decreases the stickiness of beliefs about state, thereby improving state inference. We further found that increased obsessionality, as measured by the OCI-R obsessing subscale, was associated with increased stickiness of beliefs about state and worse state inference. These findings support our second theoretical proposal, which is that obsessions result from sticky beliefs about state. As we had also hypothesized, escitalopram and obsessions had opposing effects, both on belief stickiness (decreasing and increasing belief stickiness, respectively) and on state inference (facilitating and hindering state inference, respectively). These mirror effects may explain the therapeutic effect of SSRIs on OCD^52,82,83^: they may decrease the belief stickiness that underlies obsessions.

Our theory that serotonin in OFC decreases the stickiness of beliefs about state explains why (1) serotonin depletions^1,26,27^, particularly in OFC^2,35,75^, hamper reversal learning, (2) increasing serotonin pharmacologically improves reversal learning^3^, with that improvement being opposed by blocking serotonergic transmission in OFC ^84^, and (3) naturally occurring variation in serotonergic neurotransmission in OFC correlates with reversal-learning performance^31,32,36^. This theory also explains other aspects of cognitive flexibility beyond reversal learning, in which orbitofrontal serotonin has also been implicated. For example, serotonin depletion in the OFC of monkeys seems to produce perseverative responding in extinction^85^. Extinction involves a state change, from acquisition to extinction^10,12^, so state stickiness due to orbitofrontal serotonin depletion naturally explains such perseverative behavior.

Our finding that only sufficiently high escitalopram plasma levels reduced belief stickiness and improved state inference is consistent with the dose-dependent effects of acute SSRI administration. As in our findings, only a sufficiently high citalopram dose improved reversal learning in rats; a lower dose even worsened reversal learning^86^. Acute SSRI administration increases serotonin in the raphe, which binds to autoreceptors, decreasing serotonin-neuron firing and therefore serotonin efflux^76,87,88^. Whether acute SSRI administration increases^89^ or even decreases^90^ serotonin in target regions depends, therefore, on whether the postsynaptic serotonin transporter (SERT) blockade is sufficient to more than compensate for the reduced serotonin efflux. This balance depends on dose^76^; acute citalopram administration, for example, increases serotonin in frontal cortex at higher, but not lower, doses^91^. We did not manipulate SSRI dose, but we measured plasma concentration, which, like dose, relates monotonically to SERT occupancy^92,93^. Our findings are consistent with the idea that only sufficiently high SERT occupancy increases serotonin in frontal cortex (including the OFC), thereby reducing belief stickiness and improving state inference. This idea may also explain why some studies found no differences in perseverative errors following reversal learning in participants receiving acute citalopram^94^ or escitalopram^95^ relative to placebo. Had these studies measured plasma drug levels, they could conceivably have found that sufficiently high drug plasma levels reduced those errors. Indeed, SSRIs, administered acutely at sufficiently high doses or chronically, improve reversal learning in animals^3,86^.

Chronic SSRI treatment more unequivocally increases serotonin in target regions^96^, presumably because of autoreceptor desensitization^87^. Chronic treatment should therefore further reduce belief stickiness and improve state inference.

Our theory was motivated by the hypothesized role of serotonin in modulating attractor states in the OFC^33^; it is not intended to encompass serotonin’s multiple functions, possibly mediated by other regions. Many studies^1,2,32,35,36^ (albeit not all^75^) suggest that serotonin in OFC specifically affects performance following reversals in reversal learning, whereas serotonin in other regions also affects performance before the first reversal^35,36^. Of course, our pharmacological manipulation was systemic, so we also investigated whether escitalopram modulated two other putative serotonin functions that could have played a role in our task: the Go bias^78^ and punishment^79–81^, including punishment-induced inhibition of a previously rewarded response^2,28^. We found no evidence that escitalopram modulated these functions. Thus, at least in our task, escitalopram’s effect on belief stickiness predominated over these other possible effects.

We were unsurprised that escitalopram did not modulate the inhibition of a previously rewarded response when it subsequently leads to punishment. The findings relating serotonin to that process were obtained with reversal-learning tasks^2,28^, so they are confounded by state stickiness: inhibiting a previously rewarded response when it starts to lead to punishment depends on inferring a state change. We were more surprised that escitalopram did not modulate punishment more generally or the Go bias—which we hypothesize might be mediated by serotonergic projections to regions other than the OFC. Other studies have found that a single dose of citalopram^94^ or escitalopram^95^ impaired performance in the initial acquisition of a reversal-learning task in humans, which suggests that the systemic effects of an acute SSRI dose extend beyond belief stickiness (presumably to S-R-learning processes). Various other lines of evidence suggest that serotonin affects punishment^81,97^ and behavioral inhibition^97,98^. Our task was not designed primarily to test these S-R-learning effects, so our null findings for punishment and the Go bias could conceivably be due to lack of power—although our detailed analyses of their effect sizes, together with our replication of these findings in model-based and classical-statistical analyses, suggest otherwise.

Several studies have related serotonin, OCD, and obsessive-compulsive symptoms to stimulus stickiness^72–75^. Those studies used reversal-learning tasks, but analyzed behavior with S-R-learning models without state inference^73,75^. Our findings, like prior ones^69,70^, show that participants perform state inference in these tasks. Using a model without state inference to analyze these tasks can misleadingly capture in the parameters that were used effects that are due to alterations in state inference. Stimulus stickiness is especially confounded by state stickiness: stimulus stickiness refers to the tendency to continue to choose the same stimulus^73,75^; state stickiness will cause a tendency to continue to perform the response that was adequate in the previous state, which, in a model without state inference, will masquerade as a tendency to continue to choose the same stimulus. The original study relating serotonin to stimulus stickiness reported that monkeys with orbitofrontal serotonin depletions tended to persevere in their choices over longer periods^75^—as expected if, as our theory suggests, those depletions increased state stickiness. Another study found that acute tryptophan depletion increased stimulus stickiness^73^—as also expected if that depletion increased state stickiness. More broadly, the prior findings relating serotonin, OCD, and obsessive-compulsive symptoms to stimulus stickiness seem strongly confounded by state stickiness. We suggest these prior data be reanalyzed by comparing models that include state inference (and that capture state stickiness) with stimulus-stickiness models, like we did here.

Another theory suggests that serotonin modulates learning rates on the basis of uncertainty^9^. As noted above, the increase in performance from phase 2 to phase 4 is predicted by the state-inference account but not by a modulation of learning rates by volatility or uncertainty (Fig. 1d). Similarly, our finding that this increase in performance correlates with escitalopram plasma level (*Supplementary results – Replication of the positive association between escitalopram plasma level and state inference – Escitalopram plasma levels related positively to the increase in performance from phase 2 to phase 4*) is predicted by our theory that serotonin improves state inference but not by the theory that serotonin simply modulates learning rates on the basis of uncertainty.

As noted in the Introduction, we studied participants from the general population, rather than patients with OCD, because we wanted to investigate obsessionality on a continuum applicable to the general population. Still, as expected, due to the usefulness of such dimensional studies for understanding OCD^58^, our findings relate to multiple aspects of the literature on OCD. As noted in the Introduction, our theoretical idea that obsessions correspond to sticky beliefs about state relates closely to computational theories ascribing obsessions and OCD to overly strong attractors^33,47,48^. Our ideas and findings concerning obsessions are also consistent with the suggestion that OCD involves disturbances in “complex reasoning systems” that deal with complex task representations, including hidden states^99^.

A different theoretical proposal is that OCD involves increased uncertainty regarding state changes, formalized as an increased (prior) belief that states tend to change^100,101^. Our findings contradict this idea: increased stickiness of beliefs about state, cast in the language of prior beliefs about state changes, would correspond to an increased prior belief that states tend *not* to change. Our theoretical idea that obsessions correspond to sticky beliefs about state is supported not only by our findings but also by the very definition of obsessions as “recurrent and persistent thoughts, urges, or impulses”^102^: “persistent” and “recurrent” suggest stickiness and overly strong attractors^33,47,48^, not excessive state changes.

Other work has reported that patients with OCD have decreased stimulus stickiness in a reversal-learning task^72,74^. The models used in that work, however, were variants of S-R learning, without state inference, which, as discussed above, may be problematic.

When considering the application of our findings to OCD, it is important to note that, as mentioned in the Introduction, our use of “belief” is not intended to denote a declarative, explicit belief. Patients with OCD have varying degrees of insight that their obsessions are unreasonable^103,104^, so “believing” in the obsessions, declaratively, is not necessary for OCD. We use “belief” in a Bayesian sense, derived from our computational model. Activity patterns in the OFC may represent probability distributions over states^23,37^; the Bayesian beliefs represent these probabilities. These probabilities are implicit in the activity patterns of the network^37^; they need not correspond to explicit beliefs. Patients with OCD may, in fact, form accurate declarative beliefs, even in environments with sudden contingency changes, but fail to use those declarative beliefs to control behavior^105^.

We did not find any relation between obsessions and the S-R parameters. Thus, obsessions may be selectively associated with belief stickiness.

In summary, in this article we proposed and tested two theoretical ideas: one concerning serotonin and the other concerning obsessions. The first theoretical idea is that serotonin fosters cognitive flexibility by decreasing the stickiness of beliefs about state. We demonstrated this effect by showing that the SSRI escitalopram decreased such stickiness, thereby improving state inference. Specifically, sufficiently high plasma escitalopram levels reduced belief stickiness and improved state inference relative to placebo. Moreover, higher escitalopram plasma levels were associated with both greater reductions in belief stickiness and greater improvements in state inference. The second theoretical idea is that obsessions correspond to excessively sticky beliefs about state. Supporting this idea, we found that greater obsessionality was associated with stickier beliefs about state (and therefore with worse state inference). Our two theoretical proposals are largely independent, with each standing on its own merits. Considered together, however, they have the additional advantage of explaining why SSRIs decrease obsessions: obsessions correspond to sticky beliefs about state, and SSRIs decrease such stickiness.

## Supporting information

Supplementary results

## METHODS

### Participants and recruitment

Fifty right-handed healthy men participated in the study. Data were collected at the Translational Neuromodeling Unit in Zurich, Switzerland. The study protocol was approved by the local ethics committee (Cantonale Ethics Comission, KEK Nr. 2014-0514, BASEC Nr. PB 2016-01208) and was registered in the Swiss Federal Complementary Database (Kofam). All participants had normal or corrected-to-normal vision and provided written informed consent. Additional inclusion criteria were age between 18–40 years and right handedness. The exclusion criteria were current nicotine use of more than 10 cigarettes a week, current use of recreational drugs, current severe medical condition, current medication usage, current tachycardia > 100bpm or cardiac arrhythmia, implants or conditions that would prevent a measurement in a magnetic resonance imaging (MRI) scanner, allergy to lactose or escitalopram, and past or present neurological, psychiatric, kidney, liver, or cardiac disorders. Participants were screened prior to the first experimental session to assess their general health (including the acquisition of an electrocardiogram).

### Experimental protocol

This study was part of a larger protocol using a randomized, double-blind, placebo-controlled, within-subject crossover design. Participants took part in two identical experimental sessions on different days. Each participant was administered a single-dose capsule of escitalopram (trade name: Cipralex®; dose: 15 mg; plasma peak level: 2-4 hr; half-life: 27-33 hr; bioavailability: 80%) in one session, and an identical-looking placebo (lactose), with the same shape and weight, in the other session. Half of the participants were administered escitalopram in the first session and the other half in the second session. Behavioral and brain-imaging data were acquired starting approximately 2.5 hr after capsule administration. Specifically, participants performed two cognitive tasks in a Philips 3T Ingenia scanner at the Laboratory for Social and Neural Systems, Zurich. In this article, we focus only on the behavioral data from one of those tasks: a novel Go/NoGo reversal-learning task that we call the shell task. As this task was a learning task, we restricted our analyses to the first experimental session, to avoid confounds due to prior learning that could be carried from the first to the second experimental session. Given that we only analyze the data from the first session, for simplicity we refer to participants’ drug status in terms of their status in the first session (e.g., when we talk about the “on-escitalopram group,” we are referring to the group of participants who received escitalopram in the first session).

### Calculation of escitalopram plasma levels

Escitalopram plasma levels were measured by liquid chromatography – tandem mass spectrometry (LC-MS/MS). The method is fully validated and accredited according to ISO 17025. Precision of the method is < 7.1%. Plasma levels were measured before and after the execution of the two cognitive tasks; the levels used for data analysis were calculated for the onset time of the shell task through linear interpolation of these two measurements. When the measurements yielded plasma levels below 5.0 nmol/l, which corresponds to the lower limit of quantification, we used a value of 2.5 nmol/l instead (corresponding to the average between 0 and 5.0 nmol/l); this was the case for three participants.

### Task

The shell task is a novel Go/NoGo reversal-learning task that we developed to assess state inference and belief stickiness. During the task, participants were repeatedly presented with one of six different colored and patterned shells. On each trial, they could decide to either collect the presented shell to obtain its content (Go) or to let it pass (NoGo; Fig. 1a). To collect a shell—that is, to perform Go—participants had to press the button on the side on which the bucket appeared on screen (left or right), which changed randomly across trials. To perform NoGo, participants simply had to withhold from responding during a time window of 750ms. When a shell was collected, it was cracked open, showing either a pearl (reward, +1 point), dirt (punishment, -1 point), or nothing (neutral outcome, 0 points; Fig. 1a). Crucially, different shells went through different seasons (states), in which they were more likely to contain rewards, punishments, or neutral outcomes (Fig. 1b). Each of the six shells went through two seasons twice. The season pattern of each shell was unique (Fig. 1c). For instance, the rewarding-neutral-rewarding-neutral shell started in a rewarding season (80% pearls, 20% empty) and then alternated with a neutral season (100% empty), while the punishing-neutral-punishing-neutral shell started in a punishing season (80% dirt, 20% empty) and then alternated with a neutral season (100% empty). Seasons varied independently across shells. To minimize order effects, every participant played through a different, pseudo-randomized trace of 216 trials (Fig. 1e). Rewarding and punishing seasons lasted 7-13 repetitions per shell (mean = 10), whereas neutral seasons lasted 5-9 repetitions per shell (mean = 7). Neutral seasons were shorter because they were deterministic, whereas rewarding and punishing seasons were probabilistic (Fig. 1b). All participant-specific trial sequences were generated such that the rewarding-punishing-rewarding-punishing and punishing-rewarding-punishing-rewarding shells contained 40 trials each; the remaining shells, all of which included neutral seasons, contained 34 trials each (because, as noted above, neutral seasons were shorter). Each sequence had a total of 80 rewarding trials, 80 punishing trials, and 56 neutral trials. No more than two neutral outcomes in a row were presented for a given shell during a rewarding or punishing season to avoid confusion with a neutral season.

Participants were instructed to collect as many pearls as possible while avoiding collecting dirt. They were informed that “each shell can go through different phases, in which it is more or less likely to produce certain outcomes” and were told to think of those phases as “shell seasons”; they were also explicitly told that “[d]ifferent shells may change season at different times” (Methods Fig. 1). Participants were not told how many seasons there were, the patterns of season changes, or the outcome probabilities associated with each season; all these had to be learned over the course of the task. To incentivize attentive task play, participants were informed that they would receive performance-based monetary compensation at the end of the task. This compensation was calculated on the basis of the difference between collected pearls and dirt.

Prior to the main task, participants went through a training period in which they first went through guided instructions that showed them how to collect or not collect a shell (2 trials). They then played on their own through a series of 15 training trials with 3 shells different from those used in the main task. The outcome probabilities for those shells were different from those used for any shell during the main task, to avoid knowledge transfer from the training period to the main task.

### Questionnaires

#### Obsessive-Compulsive Inventory-Revised (OCI-R)

As part of the broader protocol in which this study was included, participants filled out several questionnaires. In this study, we analyzed only the OCI-R^55^ questionnaire (which was administered in its German version^106^) because our *a priori* hypotheses were specifically related to obsessions. The OCI-R is an 18-item questionnaire that was designed as a self-report instrument to determine the diagnosis and severity of OCD, but which is sufficiently sensitive to allow the discrimination of different severities of subclinical obsessive-compulsive traits in healthy individuals^55,107^. The OCI-R is a shorter, revised version of the original OCI^108^.

#### Debriefing questionnaire

After completion of the shell task, participants filled out a debriefing questionnaire. In that questionnaire, participants were asked, among other questions, if they had understood and recognized that there were “good” and “bad” shell seasons. Participants who answered this question negatively were considered to not have understood the task and were therefore excluded from all subsequent data analyses. Participants were also considered to not have understood the task, and were therefore also excluded from data analyses, if they performed the same response (Go or NoGo) in more than 90% of the trials.

**Methods Fig. 1.**
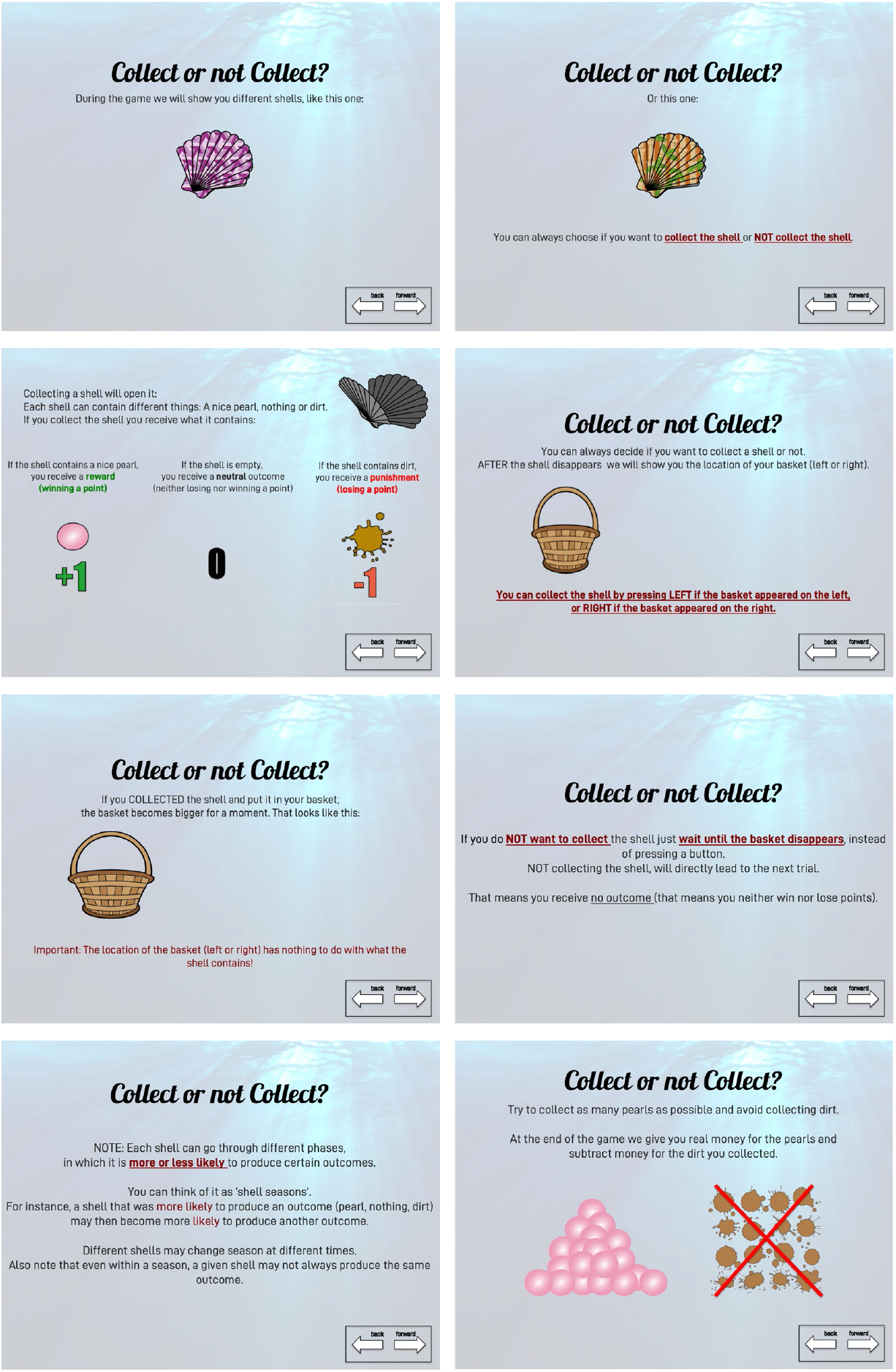
Instructions of the shell task. Participants used a response box to move to the next (or previous) slide. In the figure, we show the slides in English, ordered from top left to bottom right; in the experiment, the slides were in German and were shown one at a time. The training period started after the last instruction slide.

### Computational models

As noted in the main text, computational models belonged to two families: S-R and S-S-R (Fig. 2; Methods Fig. 2).

#### Q learning

All models from both families used (variations of) *Q* learning for action learning and selection^63,71^. *Q* learning involves learning *Q* values, *Q*(***s***, *a*), for pairs of stimuli or states, ***s***, and actions, *a*. We represent ***s*** in bold because, in our case, ***s*** was a vector (as explained below). The possible actions *a* were Go or NoGo.

*Q* values were learned using (variations of) the standard *Q* learning equations:

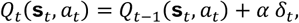

where *α* is the individual’s learning rate, and *δ*_*t*_ is a prediction error. In turn, *δ*_*t*_ was defined as:

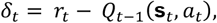

where *r*_*t*_ was –1, 0, or +1 for dirt, empty shells, and pearls, respectively. *Q* values were updated independently for each ***s***, so the subscript *t* indicates the *t*^th^ trial for state ***s***_*t*_, not the *t*^th^ trial in the task.

All *Q* values were initialized to 0. Given that no outcome was received following a NoGo action, *Q*(***s***_*t*_, NoGo) always remained equal to 0. Thus, the equations above were used only to learn *Q*(***s***_*t*_, Go).

On each trial, the model calculated the probability of performing Go using the following formula:

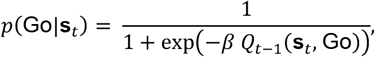

where *Q*_*t*−1_(***s***_*t*_, Go) is the previously learned *Q* value for the Go action for ***s***_*t*_, and *β*, the inverse temperature, is a free parameter that determines how much an individual’s decision is influenced by the learned *Q* value (as opposed to being random). This formula can be derived straightforwardly from the standard softmax choice rule for *Q* learning considering that *Q*(***s***_*t*_, NoGo) was always 0.

Given that the only possible actions are Go and NoGo,

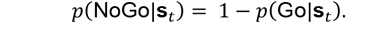

**Methods Fig. 2.**
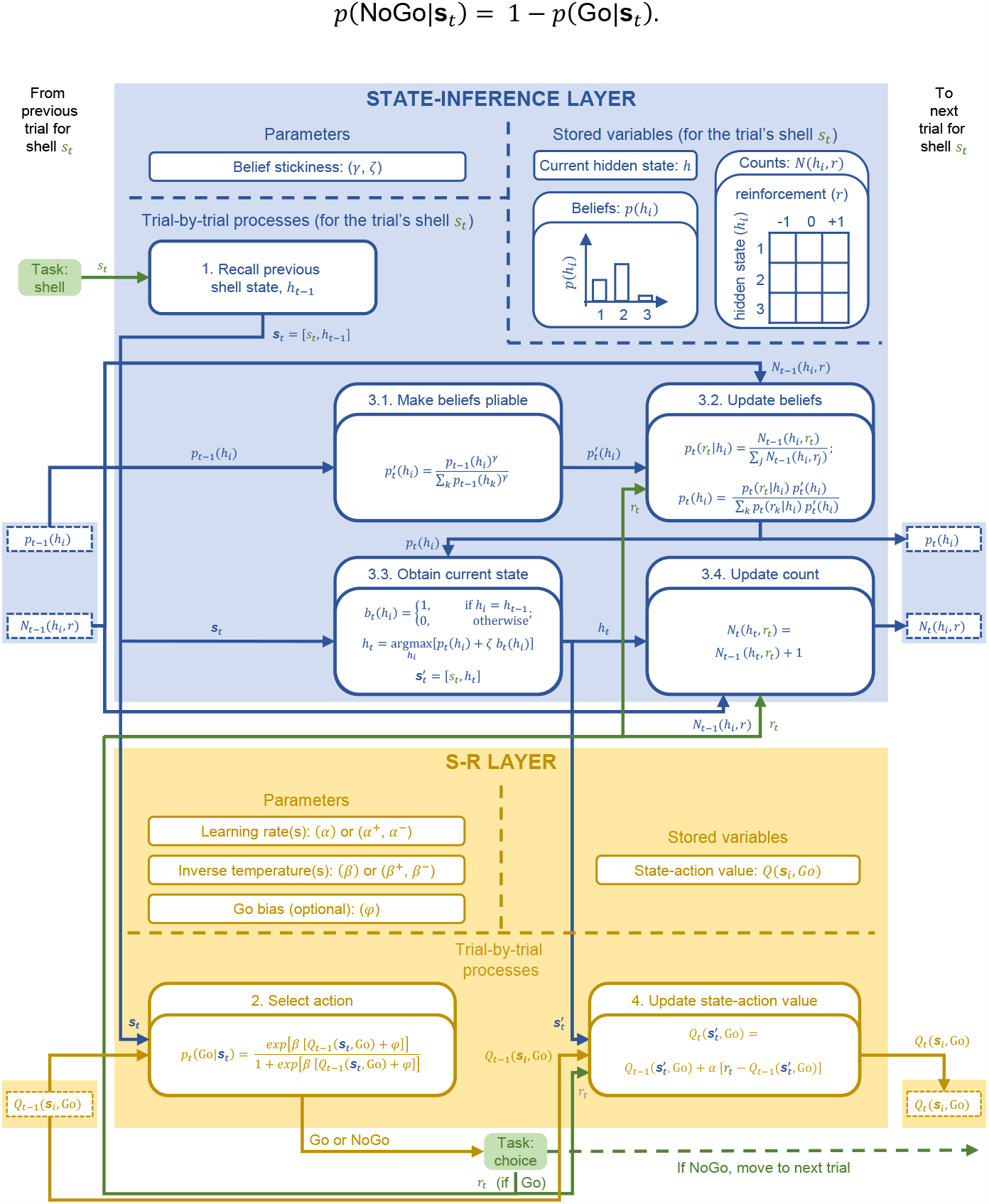
The model space was divided into two families: one implemented simple stimulus-response (S-R) learning (S-R models); the other augmented S-R learning with state inference (S-S-R models). S-R models only included the S-R layer (yellow/orange shading); S-S-R models included, in addition, the state-inference layer (blue shading). In the S-R layer, as is common in *Q*-learning^7,63^, action selection used a softmax function (box 2), and *Q* values were updated using prediction errors, corresponding to the difference between the previous *Q* value and the received reinforcement, *r*_*t*_ (box 4). *Q* values were updated only when the selected action was Go; the task did not include feedback for NoGo, so there was no *Q* learning for NoGo. S-R models and S-S-R models differed only in their treatment of the state. In S-R models, the state was univocally determined by the shell presented on the trial, *s*_*t*_. Thus, in S-R models, the state information entering boxes 2 and 4—***s***_*t*_ and 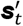, respectively—equaled just *s*_*t*_ (not shown in the figure). Given that S-R models consisted of only the S-R layer, they implemented only boxes 2 and 4. S-S-R models, in contrast, tried to infer the hidden state, *h*, of the shell presented on the trial, *s*_*t*_. Thus, in S-S-R models, the state, ***s***_*t*_, consisted of an ordered pair: ***s***_*t*_= [*s*_*t*_, *h*]. At the beginning of the trial, when the shell was presented, S-S-R models recalled the previously inferred state of the shell, *h*_*t*–1_ (box 1) and provided the pair ***s***_*t*_= [*s*_*t*_, *h*_*t*–1_] to the S-R layer, which used it for action selection (box 2). The state-inference layer then made the prior beliefs about the shell’s hidden state pliable, to allow for the possibility that the shell’s hidden state could have changed. The distribution of beliefs about the possible hidden states of the shell was represented by a categorical distribution, *p*(*h*_*i*_), over the possible hidden states, *h*_*i*_. The prior beliefs, *p*_*t*−1_(*h*_*i*_), were made pliable by flattening this categorical distribution into a new distribution, 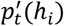, which preserved the belief ordering but made beliefs more equiprobable (box 3.1; *Methods – Mathematical proof of the properties of the formulation of belief pliability*). The extent to which the prior distribution was flattened was controlled by a belief-stickiness parameter, *γ* (0 < *γ* < 1). Larger values of this parameter corresponded to less flattening and therefore greater belief stickiness (*Methods – Mathematical proof of the properties of the formulation of belief pliability*); for example, a value of 1 would correspond to the (implicit) assumption that shells do not change state (which would go against the task instructions, which explicitly mention that shells go through different seasons). If the action selected in the trial was NoGo, there was no feedback, so there were no further changes to the state information, nor was there any learning. If the action selected was Go, however, this resulted in a reinforcement, *r*_*t*_, which was used both for state inference in the state-inference layer and for *Q* learning in the S-R layer. The state-inference layer used this reinforcement to try to infer the shell’s current state, *h*_*t*_, by using Bayesian belief updating to calculate a posterior distribution of beliefs, *p*_*t*_(*h*_*i*_), that took the observation of *r*_*t*_ into account (box 3.2). The shell’s possibly changed hidden state, *h*_*t*_, was then obtained by choosing the state with the maximum *a posteriori* belief, but with a parameter-dependent tendency to stick with the previously inferred hidden state (box 3.3). This tendency was parameterized by another belief-stickiness parameter, *ζ*. Again, larger values of *ζ* corresponded to greater belief stickiness, fostering perseveration in the belief that the shell’s hidden state remained the same, even when the evidence accumulated in the posterior beliefs, *p*_*t*_ (*h*_*i*_), indicated that it had changed. In S-S-R models, learning occurred only following these steps to obtain the shell’s current state, 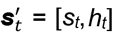, because learning applied to the currently inferred state. Thus, after obtaining this state, the state-inference layer provided it to the S-R layer, where learning then occurred, updating the *Q*-value for the shell’s newly inferred state, 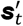 (box 4). Finally, in the state-inference layer, the count of reinforcements for the currently inferred hidden state was incremented by one for reinforcement *r*_*t*_, reflecting the fact that reinforcement *r*_*t*_ was observed in the trial (box 3.4). [The state-inference layer kept a table, *N*(*h*_*i*_, *r*), counting how many times each of the reinforcements (−1, 0, and +1) had occurred for each hidden state.] Each shell had its own replica of the “stored variables” indicated in the state-inference layer; the trial-by-trial processes in the state-inference layer used and manipulated the information specific to the shell presented in the trial.

#### S-R models

In S-R models, ***s***_*t*_ was univocally determined by the shell, *s*_*t*_, presented in the trial:

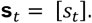

There was therefore a single *Q*_*t*_(***s***_*t*_, Go) value for each shell, which was updated whenever the participant performed Go for that shell. The differences between the eight S-R models concerned therefore how *Q*_*t*_(***s***_*t*_, Go) values were updated and used during action selection.

The simplest model, model S-R-1, used a single learning rate *α* and a single inverse temperature *β*. Model S-R-2 used two learning rates, *α*^+^ and *α*^−^, depending on whether the prediction error (*δ*_*t*_) was positive or negative, respectively. Model S-R-3 used two inverse temperatures, *β*^+^ and *β*^−^, depending on whether *Q*_*t*_(***s***_*t*_, *Go*) was positive or negative, respectively. Model S-R-4 used both two learning rates (*α*^+^ and *α*^−^) and two inverse temperatures (*β*^+^ and *β*^−^). The consideration of different learning rates, depending on the sign of *δ*_*t*_, and different inverse temperatures, depending on the sign of *Q*_*t*_(***s***_*t*_, Go), was motivated by the differential contributions of the direct and indirect pathways through the basal ganglia during action learning and selection^6,61,63,109^. Models S-R-5 through S-R-8 were equivalent to models S-R-1 through S-R-4, respectively, but included an additional parameter, *φ*, that captured a possible general bias for performing a Go response^62^. Specifically, *φ* increased the tendency to perform Go by being added to *Q*_*t*−1_(***s***_*t*_, Go) in the choice rule:

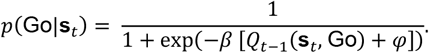

Parameter *φ* captures both motor impulsivity and the propensity for information seeking (given that only Go responses led to outcomes).

The most complex S-R model, S-R-8, thus had five free parameters (*α*^+^, *α* ^−^, *β*^+^, *β*^−^, *φ*) and obeyed the following equations:

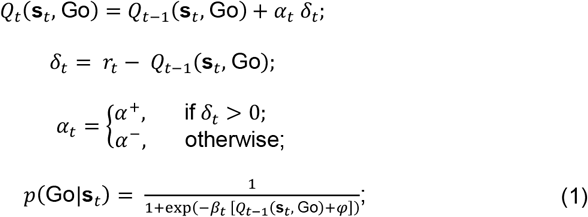

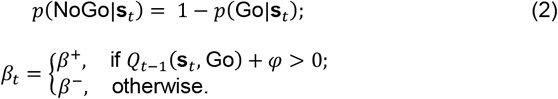

Models S-R-1 through S-R-7 were special cases of this more general model, in which, depending on the specific model, one or more of the following restrictions were applied to the more general model: *α*^+^ = *α* ^−^ = *α*, in models S-R-1, S-R-3, S-R-5, and S-R-7; *β*^+^ = *β*^−^ = *β*, in models S-R-1, S-R-2, S-R-5, and S-R-6; and *φ* = 0, in models S-R-1 through S-R-4 (Methods Table 1).

**Methods Table 1.** Parameterization of S-R models. Models varied in terms of whether they used a single learning rate (*α*) or two learning rates (*α*^+^, *α*^−^), a single inverse temperature (*β*) or two inverse temperatures (*β*^+^, *β*^−^), and whether they included a Go bias (*φ*) or not (*φ* = 0). The family of S-R models encompassed all possible eight combinations of these parameterizations, defining models S-R-1 through S-R-8. The checkmarks indicate the parameterization of each model. Models S-R-1 and S-R-8 were the simplest and most complex models, respectively.

| Model | Learning Rate(s) |  | Inverse Temperature(s) |  | Go Bias |  |
| --- | --- | --- | --- | --- | --- | --- |
| | $\alpha$ | $\alpha^+, \alpha^-$ | $\beta$ | $\beta^+, \beta^-$ | $\varphi$ | $\varphi = 0$ |
| S-R-1 | ✓ |  | ✓ |  |  | ✓ |
| S-R-2 |  | ✓ | ✓ |  |  | ✓ |
| S-R-3 | ✓ |  |  | ✓ |  | ✓ |
| S-R-4 |  | ✓ |  | ✓ |  | ✓ |
| S-R-5 | ✓ |  | ✓ |  | ✓ |  |
| S-R-6 |  | ✓ | ✓ |  | ✓ |  |
| S-R-7 | ✓ |  |  | ✓ | ✓ |  |
| S-R-8 |  | ✓ |  | ✓ | ✓ |  |

#### S-S-R models

##### State representation and interaction with the S-R layer

Unlike in S-R models, in S-S-R models, the shell presented in the trial, *s*_*t*_, did not univocally determine ***s***_*t*_. Instead, S-S-R models tried to infer the shell’s hidden state, *h* (i.e., its underlying season), using a state-inference layer placed atop the S-R learning layer (Fig. 2; Methods Fig. 2). The state representation in S-S-R models therefore consisted of a pair shell-hidden state, ***s*** = [*s, h*]. There were eight S-S-R models (S-S-R-1 through S-S-R-8), each of which added the aforementioned state-inference layer to the corresponding S-R model (S-R-1 through S-R-8, respectively).

At the beginning of a trial, when presented with the shell, *s*_*t*_, the state-inference layer in S-S-R models started by recalling the previous shell state, *h*_*t*−1_ (box 1 in Methods Fig. 2).

It then provided the pair ***s***_*t*_ = [*s*_*t*_, *h*_*t*−1_] to the S-R layer, which used ***s***_*t*_ as the state information for action selection [equation (1); box 2 in Methods Fig. 2]. Thus, in S-S-R models, actions were selected on the basis of the *Q* values learned for the specific shell-hidden state pair.

If the action selected was NoGo, there was no feedback, so there was no new information about the shell’s state and nothing to learn; thus, the state-inference layer just implemented a step aimed at making beliefs flexible (box 3.1 in Methods Fig. 2), which we describe below. If, however, the action selected was Go, that led to a reinforcement, *r*_*t*_, which both provided new information about the shell’s hidden state and supported new learning about the *Q* value for the (newly inferred) shell state. Thus, following a Go, in addition to the aforementioned step aimed at making beliefs flexible (box 3.1 in Methods Fig. 2), the state-inference layer tried to infer the shell’s current state, *h*_*t*_, and updated the relevant state variables (boxes 3.2–3.4 in Methods Fig. 2), using steps that we detail below. It then provided the pair 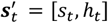 to the S-R layer, which updated the *Q* value specifically for this 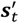, which already included the shell’s newly inferred state (box 4 in Methods Fig. 2).

##### State inference

For each shell, the state-inference layer stored three variables: a categorical probability distribution representing the current beliefs that the shell is in each of various possible hidden states, *p*(*h*_*i*_); the specific hidden state that the state-inference layer believes the shell is currently in, *h* [which is partly, but not solely, determined by *p*(*h*_*i*_), as described below]; and a tabulation of how many of each of the three types of reinforcement (–1, 0, and +1) were received in each hidden state, *N*(*h*_*i*_, *r*), where *r* ranges over –1, 0, and +1 (Methods Fig. 2). The role of the state-inference layer was to keep these variables up to date, to always be able to best infer the shell’s current hidden state. This process, however, was modulated by two belief-stickiness parameters, *γ* and *ζ*. If these parameters were high, beliefs about the states became excessively sticky (i.e., resistant to change), making state changes difficult to detect; in other words, the state-inference layer became “stuck” in previously acquired beliefs, ignoring new, incoming evidence to the contrary.

State inference consisted of four steps (boxes 3.1–3.4 in Methods Fig. 2). The first step, applied at the beginning of the trial, to avoid getting stuck in the previously inferred beliefs, was to make the previous beliefs pliable, for the shell presented (box 3.1 in Methods Fig. 2). The task instructions explicitly noted that shells go through different seasons. Thus, while the previous beliefs encoded in the categorical probability distribution over hidden states, *p*_*t*−1_(*h*_*i*_), reflected the previous best knowledge about the most probable hidden state(s) for the shell, that knowledge might no longer be current because the shell’s season could have changed. A flexible learner would allow for the possibility that the previously inferred beliefs might no longer be current. An inflexible learner, characterized by excessive belief stickiness, in contrast, would stick more rigidly to the previously inferred beliefs. To allow the model to consider the possibility that the previously inferred beliefs might no longer be current—thereby allowing it to more flexibly infer that the state has changed when there was evidence for such a change—we made beliefs about the shell state pliable, by drifting them in the direction of the uniform distribution. To be able to characterize the degree to which participants did or did not exhibit such belief flexibility (or its converse, belief stickiness), we modulated the degree of such drift by a belief-stickiness parameter *γ* (0 < *γ* < 1) that was fitted individually for each participant. We formalized this process through the following equation:

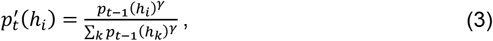

where *k* ranged over all shell states. This equation first down-weights the prior beliefs, by raising them to *γ* (0 < *γ* < 1), and then normalizes them to ensure that the sum of beliefs across all possible shell states always equaled 1.

We chose this mathematical formulation because, with 0 < *γ* < 1, it ensured that: (1) the order of the beliefs was preserved; (2) the largest belief was decreased (except if all beliefs were equal), thereby allowing for belief flexibility; (3) belief flexibility (or its converse, belief stickiness) was modulated in a parametric fashion, with larger values of *γ* corresponding to more belief stickiness—i.e., an exaggerated reliance on the previously inferred beliefs relative to the new information, thereby ignoring the possibility that the shell’s season could have changed, which resulted in decreased task performance (see *Supplementary results – Effects of γ and ζ on task performance*); and (4) the transformed belief values, like the previous belief values, obeyed the properties of a categorical distribution (Methods Fig. 3; see subsection *Mathematical proof of the properties of the formulation of belief pliability* for proof of these 4 properties). We set an upper bound of 1 for *γ* because a value larger than 1 would mean that the certainty about the shell’s state would increase—i.e., the highest value of *p*_*t*−1_(*h*_*i*_) would become even larger in 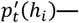— following NoGo’s, in which there is no new information about the shell state.

The preceding step (box 3.1 in Methods Fig. 2) was applied regardless of whether the action selected in the trial was Go or NoGo. The remaining steps (boxes 3.2–3.4 in Methods Fig. 2), however, were applied only when the action was a Go, as only in that case was there new information about the shell state (in the form of the received reinforcement, *r*_*t*_) that needed to be incorporated for state inference. The first of those remaining steps (box 3.2 in Methods Fig. 2) was to use *r*_*t*_ to update the categorical probability distribution over hidden states (box 3.2 in Methods Fig. 2). This update was Bayesian. First, we calculated the likelihood, *p*_*t*_(*r*_*t*_|*h*_*i*_), of observing that reinforcement *r*_*t*_ for each hidden state, *h*_*i*_, by dividing the number of times that we observed that reinforcement in that state by the total number of reinforcements that we observed in that state:

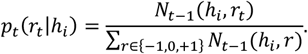

Next, we calculated the posterior for *p*_*t*_(*h*_*i*_) using Bayes rule, considering both the likelihood, *p*_*t*_(*r*_*t*_|*h*_*i*_), and the previous beliefs about *p*_*t*−1_(*h*_*i*_), already drifted towards the uniform distribution through equation (3) [hence, 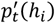]:

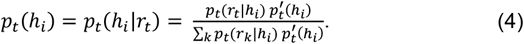

After obtaining the posterior probability distribution over hidden states, *p*_*t*_(*h*_*i*_), the next step was to obtain the shell’s current state (box 3.3 in Methods Fig. 2). In principle, the shell’s current state could be obtained as the maximum *a posteriori* (MAP) value of that posterior distribution:

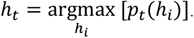

Given our focus on belief stickiness, however, we sought to characterize parametrically the possible propensity of each participant to stick with the previously inferred state of a shell, *h*_*t*−1_, even when the posterior distribution, *p*_*t*_(*h*_*i*_), suggested that there was a change in shell state. We formalized this propensity by adding *ζ*, a free parameter (0 < *ζ* < 0.5) fitted individually for each participant, for the hidden state that corresponded to the previously inferred hidden state, *h*_*t*−1_, before computing the argmax:

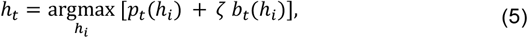

where

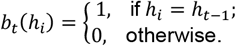

We made *ζ* < 0.5 rather *ζ* < 1 because, in practice, values of *ζ* between 0.5 and 1 would produce the same behavior, thereby causing problems with model identifiability (see *Supplementary results – Effects of γ and ζ on task performance*). Parameter *ζ* was our second belief-stickiness parameter: larger values of *ζ* caused participants to stick with the previously inferred state more, even when incoming evidence suggested that the state had changed.

After the shell’s current state, *h*_*t*_, was obtained through the preceding equations, the state layer then provided the updated state information, 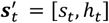, to the S-R layer. As noted above, the S-R layer then updated the *Q* value specifically for this 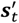, wh ich al ready includes the shell’s newly inferred state (box 4 in Methods Fig. 2).

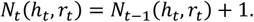

The state-inference layer then had to perform just one final step (box 3.4 in Methods Fig. 2): adding one to the count of the number of times that the obtained reinforcement, *r*_*t*_, was observed in the currently inferred state, *h*_*t*_ :

##### Variable initialization

To finish defining the S-S-R models, we need to specify how variables were initialized. In the task, but unbeknownst to participants, each shell would only alternate between two possible states (seasons), and there were three different seasons (rewarding, neutral, punishing) across shells (Fig. 1). We therefore assumed for all models that there were no more than three hidden states (seasons) per shell. Accordingly, we initialized the beliefs over the possible states for each shell, *p*_0_(*h*_*i*_), for *i* = 1 to 3, to 1/3. We initialized the count of the number of reinforcements of each type for each possible hidden state of each shell, *N*(*h*_*i*_, *r*), for *r* ∈ {−1,0, +1} to 1, which is equivalent to starting with equal probabilities of observing each reinforcement in each state. This initialization mimicked the task instructions, which only stated that shells could yield three distinct reinforcements. We initialized the current hidden state for each shell, *h*, to 1. This choice was arbitrary, as all states were otherwise equal in the beginning, so any state could be considered the initial hidden state.

Our assumption that there were no more than three hidden states was, of course, a simplification. We could instead have considered models that create states as needed^10,12,64,65,110^. We wanted to keep our models as simple as possible, however, to ensure parameter identifiability; we also did not want to create a proliferation of different model families that would hamper model selection. We therefore created models focused specifically on our theoretical hypotheses; given that these hypotheses related specifically to belief stickiness, we created models that modulated belief stickiness parametrically. We deliberately chose not to model the creation of an arbitrary number of states—or, for example, the learning of a prior that indicates when a reversal is likely to occur^111,112^—to keep the models simple and the model space manageable.

##### Mathematical proof of the properties of the formulation of belief pliability

Our task was designed to tap into belief flexibility (or its converse, belief stickiness). The task instructions specifically stated that shell seasons would change over time; thus, belief stickiness, in the form of overreliance on previously learned beliefs, would be detrimental for performance. As noted above, we made beliefs pliable between consecutive presentations of a shell by drifting the previous beliefs towards a uniform distribution, using equation (3), which we reproduce here for ease of reference:

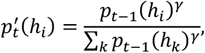

where *p*_*t*−1_(*h*_*i*_) are the previous beliefs, *k* ranges over the different shell states (from 1 to 3, as we modeled three states per shell), *γ* controls the degree of belief stickiness, and 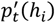 are the drifted beliefs.

As also noted above, we chose this formalization because, with 0 < *γ* < 1, it obeys four properties: (1) it preserves the order of the beliefs; (2) it decreases the largest belief, thereby allowing for belief flexibility; (3) it provides a parametric modulation of belief flexibility (or its converse, belief stickiness), with larger values of *γ* corresponding to more belief stickiness; (4) the transformed belief values obey the properties of a categorical distribution (Methods Fig. 3). Here, we prove these four properties mathematically.

For purposes of these demonstrations, we will use *b*_1_, *b*_2_, and *b*_3_ to represent the previous state beliefs, *p*_*t*−1_(*h*_*i*_), ordered from largest to smallest, such that *b*_1_ ≥ *b*_2_ ≥ *b*_3_. In other words, *b*_1_ will correspond to the largest value of *p*_*t*−1_(*h*_*i*_), *b*_2_ to the next largest, and *b*_3_ to the smallest. We can therefore write equation (3) as:

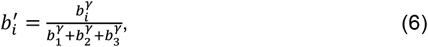

where *b*_*i*_ and 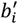 represent the previous and drifted beliefs, respectively. We will also use the following facts about our model in the demonstrations: 0 < *γ* < 1; *b*_1_ + *b*_2_ + *b*_3_ = 1; 0 < *b*_*i*_ < 1 for all *i*. We now demonstrate the four properties mathematically.

###### Property 1: Order preservation

Formally, the statement that our transformation preserves the order of beliefs corresponds to the statement that, given our ordering *b*_1_ ≥ *b*_2_ ≥ *b*_3_, then 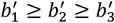. Considering equation (6), we have:

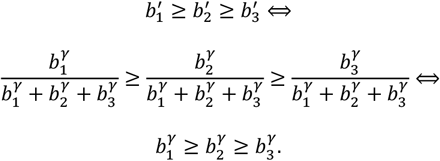

**Methods Fig. 3.**
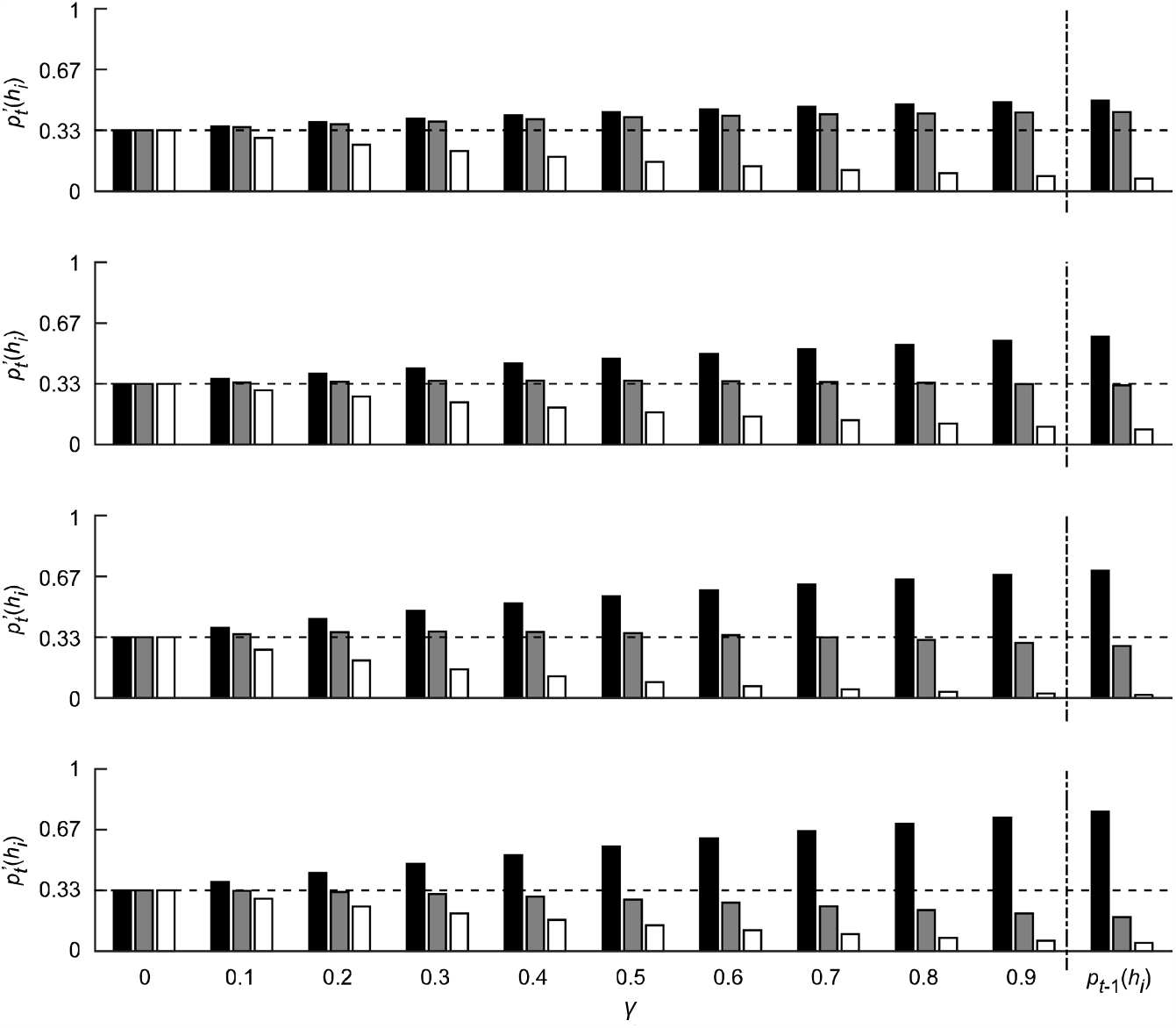
The parameter *γ* determines how pliable beliefs are—i.e., how much they drift towards the uniform distribution between consecutive presentations of the same shell. Each row represents the effects of applying equation (3) to an illustrative distribution of previous beliefs, using various levels of *γ*. The distribution of previous beliefs is shown on the right, ordered from the state with the largest belief (left bar; black) to the state with the lowest belief (right bar; white). The drift of the previous distribution following application of equation (3) is shown for a specific subset of *γ* values (from 0 to 0.9 in 0.1 increments). The larger the value of *γ*, the more similar the transformed distribution is to the previous distribution—i.e., the less the belief drift and therefore the greater the tendency to stick with the previous beliefs (which, given the nature of this task, constitutes belief stickiness). In the extreme, if *γ* were equal to 1, the previous distribution would be unchanged. The smaller the value of *γ*, the more the beliefs drift towards the uniform distribution—i.e., the more pliable beliefs become, down-weighting the previous beliefs in favor of new information. The figure illustrates visually the four properties of this transformation that make it useful for our purposes: (1) it preserves the order of the beliefs (note that, in each row, the three bars maintain their relative order for all values of *γ* > 0); (2) it decreases the largest belief (note that, in each row, the black bar is smaller for all values of *γ* than for the previous distribution); (3) it provides a parametric modulation of belief flexibility (or its converse, belief stickiness), with larger values of *γ* corresponding to more belief stickiness, as explained above; (4) the transformed belief values constitute a categorical distribution.

**Methods Fig. 4.**
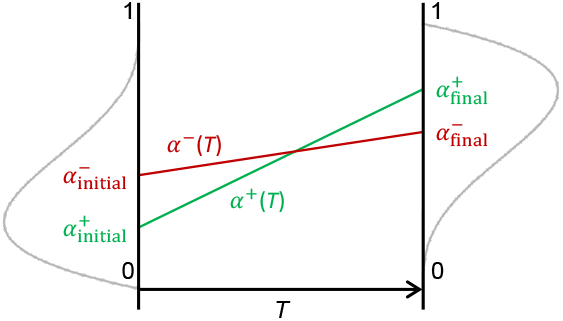
Illustration of how the positive and negative learning rates—*α*^+^(*T*) and *α*^−^(*T*), respectively—varied as a function of the overall trial in the task, *T*, in the extended S-R-8 model. Each of *α*^+^(*T*) and *α*^−^(*T*) varied linearly between an initial value (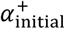 and 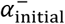, respectively) and a final value (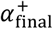 and 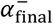, respectively). The green and red lines show an example of how *α*^+^(*T*) and *α*^−^(*T*) might vary across the task for an illustrative fictitious participant. We used beta distributions with parameters 2, 4 and 4, 2 as prior distributions for the initial (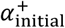 and 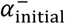) and final values (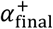 and 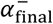), respectively. These prior distributions, illustrated with grey lines, made the final values (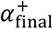 and 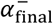) more likely to be larger than the initial values (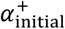 and 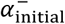): note the upward and downward shift of the prior distributions for the final and initial values, respectively, illustrated on the right and left, respectively.

Given that, in our model, *b*_*i*_ > 0 for all *i* and *γ* > 0, and that a power function with positive base and exponent increases monotonically, *b*_1_ ≥ *b*_2_ ≥ *b*_3_ implies that 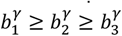, so 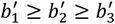.

###### Property 2: Decrease of the largest belief

Formally, the statement that our transformation decreases the largest belief corresponds to the statement that 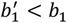. There is only one exception to this rule: if all beliefs are equiprobable, i.e., if 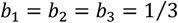, then they remain equiprobable, i.e., 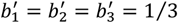. In fact, this is necessary so as not to violate Property 1.

We will first demonstrate that if there is one belief that is larger than the others, then 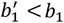. Formally, the statement that there is one belief that is larger than the others corresponds to *b*_1_ > *b*_2_ ≥ *b*_3_. Then, the statement that 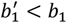corresponds to:

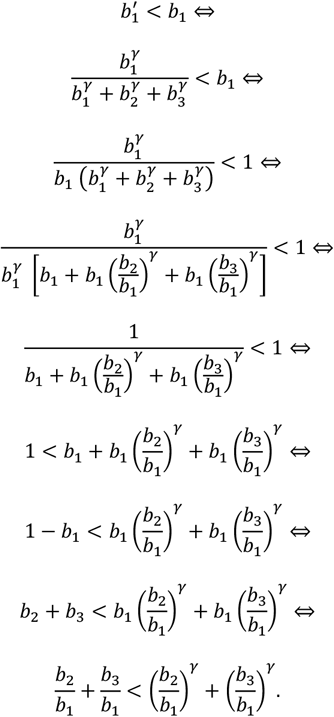

Considering the ordering *b*_1_ > *b*_2_ ≥ *b*_3_ and that *b*_*i*_ > 0 for all i, we have 0 < *b*_2_/*b*_1_ < 1 and 0 < *b*_3_/*b*_1_ < 1. Considering additionally that, in our model, 0 < *γ* < 1, we have *b*_2_/*b*_1_ < (*b*_2_/*b*_1_)^*γ*^ and *b*_3_/*b*_1_ < (*b*_3_/*b*_1_)^*γ*^. Thus, *b*_2_/*b*_1_ + *b*_3_/*b*_1_ < (*b*_2_/*b*_1_)^*γ*^ + (*b*_3_/*b*_1_)^*γ*^ is necessarily true, which shows that 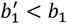.

We next consider the case in which all beliefs are equiprobable, i.e., in which *b*_*i*_ = 1/3 for all *i*. In this case, for all *i*, we have:

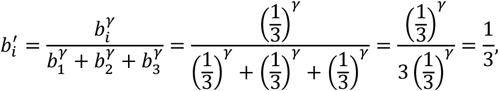

so the beliefs remain equiprobable after the transformation.

###### Property 3: Parametric modulation of belief flexibility, with larger values of γ corresponding to more belief stickiness

Given that the task instructions specifically noted that shell seasons might change, belief flexibility required the willingness to down-weight previously learned beliefs in the face of new information, whereas excessive reliance on the previously learned beliefs constituted a form of belief stickiness. Here, we show that smaller values of *γ* resulted in greater belief flexibility (and therefore, conversely, greater values of *γ* resulted in greater belief stickiness).

Belief flexibility corresponds to a down-weighting of the previous beliefs, so that new information can more easily lead to a belief change. Formally, and more specifically, then, greater belief flexibility corresponds to a greater decrease in the maximum belief—i.e., when there is one belief that is larger than the others, under our ordering *b*_1_ > *b*_2_ ≥ *b*_3_, the statement that smaller values of *γ* resulted in greater belief flexibility corresponds to the statement that the function that defines 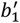 as a function of *γ* is monotonically increasing. Given equation (6), we have:

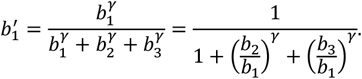

We can prove that 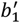 is monotonically increasing as a function of *γ* by proving that ^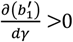^. We have:

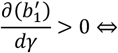

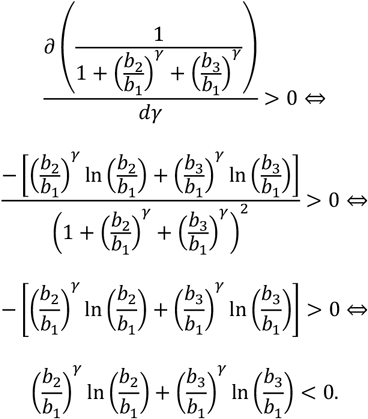

Given that *b*_*i*_ > 0 for all i, we have that (*b*_2_/*b*_1_)^*γ*^ > 0 and (*b*_3_/*b*_1_)^*γ*^ > 0. Given that *b*_1_ > *b*_2_ ≥ *b*_3_, we have that ln(*b*_2_/*b*_1_) < 0 and ln(*b*_3_/*b*_1_) < 0. Thus, (*b*_2_/*b*_1_)^*γ*^ ln(*b*_2_/*b*_1_) < 0 and (*b*_3_/*b*_1_)^*γ*^ ln(*b*_3_/*b*_1_) < 0, so (*b*_2_/*b*_1_)^*γ*^ ln(*b*_2_/*b*_1_) + (*b*_3_/*b*_1_)^*γ*^ ln(*b*_3_/*b*_1_) < 0, which in turn means that 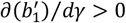, as we wanted to demonstrate.

In the special case of equiprobable beliefs, we showed above that the transformed beliefs are also equiprobable, as intended, regardless of the value of *γ*.

###### Property 4: The transformed beliefs obey the properties of a categorical distribution

To show that the transformed beliefs, 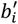, with *i* =1, 2, 3, obey the properties of a categorical distribution, we need to show that 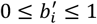 and that 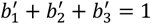. The fact that 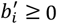 is a straightforward consequence of equation (6) together with the fact that 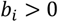 for all i. We next prove that 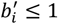:

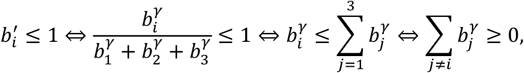

which is necessarily true given that *b*_*j*_ > 0 for all *j*. Finally, we prove that 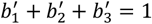:

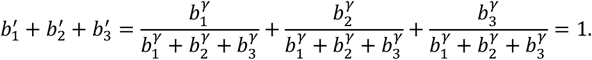

#### Alternative model families

We implemented two additional model families, designed to rule out alternative explanations for our findings.

##### S-R models with varying learning rate

As noted in the main text, state inference improves performance when states are revisited. Given that states are revisited later in the task, however, an alternative explanation for better performance when states are revisited might be that, rather than performing state inference, participants simply get faster at learning the task contingencies over time. To rule out this alternative explanation, we also implemented a family of S-R models in which participants could progressively learn faster (or slower; see below) during the task (but in which they did not perform state inference). Specifically, in this family of models, learning rates changed from trial to trial, allowing them to become faster (or slower). Like the other model families, this family consisted of eight models, each of which changed the corresponding S-R model (S-R-1 through S-R-8) to have variable learning rates.

In this family, *α*^+^ and *α*^−^ varied independently as a function of trial: *α*^+^(*T*) and *α*^−^(*T*), respectively, where *T* is the trial. (We capitalize “*T*” to indicate that it represents the overall trial in the task; elsewhere, we use lowercase “*t*” to represent the *t*^th^ trial of a specific shell.) To ensure that *α*^+^(*T*) and *α*^−^(*T*) always remained between 0 and 1, they varied linearly between an initial value (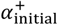 and 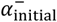, respectively) and a final value (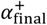 and 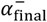, respectively), with both the initial and final values constrained to be between 0 and 1 (Methods Fig. 4). In other words, the learning rate *α*(*T*) on trial *T* was:

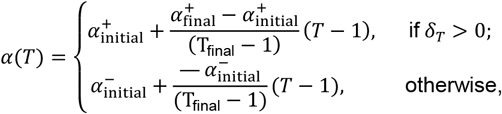

where T_final_ is the total number of trials, *δ*_*T*_ is the prediction error on trial *T*, and 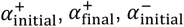, and 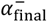 are free parameters. Given that we were specifically interested in testing the hypothesis that learning became faster across the task, we used different priors for the initial (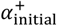 and 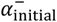) and final (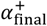 and 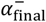) values that made it more likely that the final values would be larger than the initial values (Methods Fig. 4). Specifically, we used beta distributions with parameters 2, 4 and 4, 2 as the priors for the initial and final values, respectively. All other aspects of these models were unchanged from the base S-R models.

##### S-R models with stimulus stickiness

Several studies have related serotonin, OCD, and obsessive-compulsive symptoms to “stimulus stickiness”^72–75,113^. These studies used reversal-learning tasks consisting of a choice between two stimuli. In that context, stimulus stickiness was defined as the tendency to continue to choose the same stimulus just because it had been chosen recently. The equivalent in our task is continuing to collect a shell—i.e., do Go—just because it was collected recently.

The implementation of stimulus stickiness in the aforementioned studies has not always been uniform^72–75,113^. We used the original formulation^75,113,114^ because it allows stimulus stickiness to build up and persist across several trials. The alternative formulation considers stimulus stickiness to depend only on the previous trial^72–74^, which seems both less complete and less plausible psychologically. Moreover, the original formulation was used in a study with orbitofrontal serotonin depletion^75^, so it bears most directly on the theoretical ideas about serotonin that we advance in this article.

Let the stimulus stickiness for shell *s*_*t*_ presented on trial *t* be represented by *k*_*t*_(*s*_*t*_). In the beginning of the task, the stimulus stickiness for all shells is initialized at 0. Then, on trial *t, k*_*t*_(*s*_*t*_) is updated as follows:

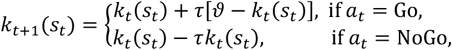

where *τ* and *ϑ* are free parameters that represent, respectively, the learning rate for stimulus stickiness and the maximum value for stimulus stickiness. Thus, if the action selected was a Go, the stimulus stickiness for the shell is increased (towards *ϑ*); if the action selected was a NoGo, the stimulus stickiness for the shell is decreased (towards 0).

During action selection, the stimulus stickiness for the shell presented in the trial increases the tendency to do Go. Thus, the softmax selection rule [equation (1)] becomes:

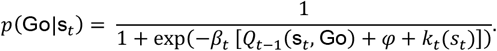

For simplicity, here we represent s_*t*_ as a scalar rather than a vector, because stimulus-stickiness models were an extension of S-R models; as noted in section *S-R models* above, in S-R models, **s**_*t*_ = [*s*_*t*_].

#### Model inversion

We implemented a total of 32 models: 8 S-R models (*Methods – Computational models – S-R models*), 8 S-S-R models (*Methods – Computational models – S-S-R models*), 8 S-R models with varying learning rates (*Methods – Computational models* – *Alternative model families – S-R models with varying learning rate*), and 8 S-R models with stimulus stickiness (*Methods – Computational models – Alternative model families – S-R models with stimulus stickiness*). We inverted each of these 32 models separately for each participant, using the participant’s choice behavior. This process yielded, for each participant–model pair, a set of parameter estimates and an approximation of the log model evidence (LME).

We calculated the log likelihood (LLH) of observing a given participant’s sequence of Go/NoGo actions, ***a***, as a function of the trial-by-trial sequence of shell presentations, ***shell***, the trial-by-trial sequence of reinforcements, ***r***, the model, *M*, and the set of free model parameters, ***θ***^115^:

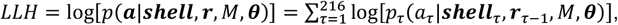

where *τ* represents the trial number, ***shell***_*τ*_ represents the sequence of shells until, and including, trial *τ*, and ***r***_*τ*−1_ represents the sequence of reinforcements until, but excluding, trial *τ*. We calculated the probability of observing each action, *a*_*τ*_, using equations (1) and (2).

We performed Bayesian inference on the parameters using a Markov chain Monte Carlo (MCMC) inversion routine based on the Metropolis-Hastings algorithm with Gibbs sampling^116^. We used truncated Gaussians for the proposal distributions, with probability density functions *p*(*x*_*i*_|*μ*_*x*_, Σ_*x*_, *LB*_*x*_, *UB*_*x*_), where *x*_*i*_ denotes a given value of the parameter of interest, *x*, and *μ*_*x*_, Σ_*x*_, *LB*_*x*_, and *UB*_*x*_ denote, respectively, the mean, covariance matrix, and lower and upper bounds of the respective distribution.

We set the prior distribution of parameters that ranged between 0 and 1 (*α, α*^+^, *α*^−^, *φ, γ*, 2*ζ*) to a Beta distribution. We parameterized the Beta distribution with both hyperparameters set to 1.01, to obtain an approximately uniform prior distribution over the open interval (0, 1). In the case of *ζ*, the Beta distribution was for 2*ζ*, so the prior for *ζ* was approximately uniform over the open interval (0, 0.5). We set the prior distribution for the inverse temperatures (*β, β*^+^, *β*^−^) to a Gamma distribution, given that the inverse temperatures can, in theory, range from 0 to +∞. We parameterized the Gamma distribution with both hyperparameters set to 2, to obtain a prior that reflected reasonable tradeoffs between exploration and exploitation; these values of the Gamma hyperparameters are relatively similar to those obtained empirically with simpler models^117^.

We implemented model inversion in MATLAB (R2015a). However, we implemented the models and their likelihood functions in C, to increase estimation speed.

#### Model inversion for model comparison

To compare models, we estimated the LME of each model for each participant’s choice sequence, ***a***, using a set of MCMC inversions with thermodynamic integration. We chose thermodynamic integration because it produces a more accurate estimate of the LME than other sampling methods^66,67^, and simpler methods, such as the Akaike information criterion or the Bayesian information criterion, do not accurately penalize model complexity^118^. We used a population MCMC approach, parallel tempering, which draws from a number of chains at different inverse temperatures—*ω*_*chain*_, varying from 0 to 1 (with no relation to the inverse temperatures *β* used in the models)—and we computed LMEs through thermodynamic integration^66^. We implemented these procedures using an earlier version of the implementation that is available in TAPAS (www.translationalneuromodeling.org/tapas)^119^. Parallel tempering and thermodynamic integration have been used previously in similar contexts^120,121^.

We ran each inversion routine for LME estimation using 50 chains with a 5-th order temperature schedule^67,122^ to increase statistical efficiency^67,123^; the inverse temperature of each chain, *ω*_*chain*_, was therefore:

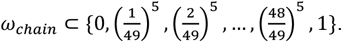

We set the maximum number of iterations per chain to 10000, and we discarded the first 5000 samples of each chain, from an initial warm-up phase^123^. We allowed up to 8 swaps between chains per iteration, to minimize the risk of an individual chain converging towards a local minimum and therefore to improve the sampling of potentially multimodal distributions^67^.

We assessed convergence of these routines by calculating the maximum relative absolute deviation between the estimated LMEs (which were calculated using 5000 iterations from 50 chains at different temperatures, as described above) and the LMEs that would have been estimated if using instead 1250, 2500, or 5000 iterations from 10 to 50 chains at distinct temperatures. We found that across all 44 subjects, 32 models, the 3 tested numbers of iterations, and the 41 chains, the maximum relative absolute variation in the estimated LME values was 1.64%, which strongly suggests that the settings used to perform the model-inversion routines were appropriate.

#### Model inversion for parameter estimation

To estimate the parameters for the winning model, we used a similar procedure. Specifically, we also used MCMC with a maximum number of 10000 iterations per chain, from which we discarded the first 5000 samples (corresponding to the warm-up phase), and in which we allowed up to 8 swaps between chains per iteration. In this case, however, we used 10 chains with an inverse temperature of 1, as we wanted all chains to converge to the same posterior distribution (rather than having a differential weighting of the prior versus the likelihood distributions from chain to chain, as in thermodynamic integration).

We assessed convergence of these routines by calculating the potential scale reduction factors, 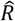, for each parameter and for the log-likelihood, for each of the 44 subjects. We verified that all those 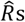 were lower than 1.1, as recommended^123^. For each model-inversion routine, we estimated the respective subject-specific parameters by averaging the non-discarded 5000 samples of the last MCMC chain—an arbitrary selection because the *R* values suggested that all chains had converged to the same distribution.

### Model comparison

To perform model comparison on the basis of the LMEs, we performed random-effects Bayesian model comparison^68,124^ using the Variational Bayesian Analysis (VBA) toolbox^125^. A model was selected as the winning model if it showed a protected exceedance probability (PEP) over 0.95^68^. The exceedance probability (EP) indicates how likely it is that a given model is more frequent, in the population studied, than any of the other models in the comparison set^124^. PEPs correct EPs for the possibility that the observed differences in model frequencies could have arisen by chance^68^.

### Statistical analyses

#### Canonical correlation analysis

All statistical analyses are described in the Results section. Here, we provide a brief description of canonical correlation analysis (CCA), for readers unfamiliar with the technique. We also explain why we chose CCA to analyze the relation of *γ* and *ζ* with escitalopram (levels and group) and OCI-R scores (OCI-R obsessing and OCI-R other).

As noted in the Results, we were interested in how the latent construct of belief stickiness, which we hypothesized underlay parameters *γ* and *ζ*, related to escitalopram (levels and group) and OCI-R scores (OCI-R obsessing and OCI-R other). CCA finds linear combinations of one set of variables (called the *y* variables) that maximally correlate with linear combinations in another set of variables (called the *x* variables). The linear combinations of both *y* and *x* variables are called canonical variates. Each canonical variate for the *y* variables is paired with a canonical variate for the *x* variables; such pairs are called canonical variate pairs. We set *γ* and *ζ* as the *y* variables, and we set escitalopram plasma level, group, OCI-R obsessing, and OCI-R other as the *x* variables.

We chose CCA for two reasons. First, CCA can find, in a data-driven manner, latent constructs in a set of variables that relate to another set of variables (or, more specifically, to linear combinations thereof); we hypothesized that CCA would find that our hypothesized latent belief-stickiness construct related to escitalopram and OCI-R (specifically, OCI-R obsessing) scores. Second, CCA naturally takes into account the correlations within each set of variables, so it handled naturally the strong correlation between *γ* and *ζ* noted in the *Results*.

We obtained *p*-values for cross-loadings in the CCA using a permutation test^126,127^.

##### Coding of group and escitalopram plasma level in regressions

Most regressions in the main text and *Supplementary results* included group (placebo vs. escitalopram) and escitalopram plasma level as independent variables (sometimes together with other independent variables). In all such regressions, we coded group and escitalopram plasma level in the same way. We coded group as 0 and 1 for participants on placebo and escitalopram, respectively. We set the escitalopram plasma level for participants in the placebo group to 0, given that these participants had, by definition, no escitalopram in their plasma. Finally, we centered the escitalopram plasma level for participants in the escitalopram group, using the mean value from participants in that group. Thus, the coefficient for group represented the difference between the escitalopram group, at its mean level of escitalopram, and the placebo group.

##### Multivariate tests

For simplicity, when several multivariate tests are available for an analysis, we report only Wilks’s Λ, as it is the most commonly reported test^128^. However, Wilks’s Λ, Hotelling-Lawley’s trace, Pillai’s trace, and, when applicable, Roy’s largest root all always produced the same conclusions as Wilks’s Λ in terms of statistical significance.

##### Software packages used for the statistical analyses

We conducted the statistical analyses in MATLAB (R2015a and R2019a), RStudio^129^ (R version 4.0.5^130^), jamovi (version 1.6.23)^131^, and IBM SPSS (Statistics 22). Methods Table 2 lists the specific functions or modules from each package that we used for each type of analysis.

**Methods Table 2.**
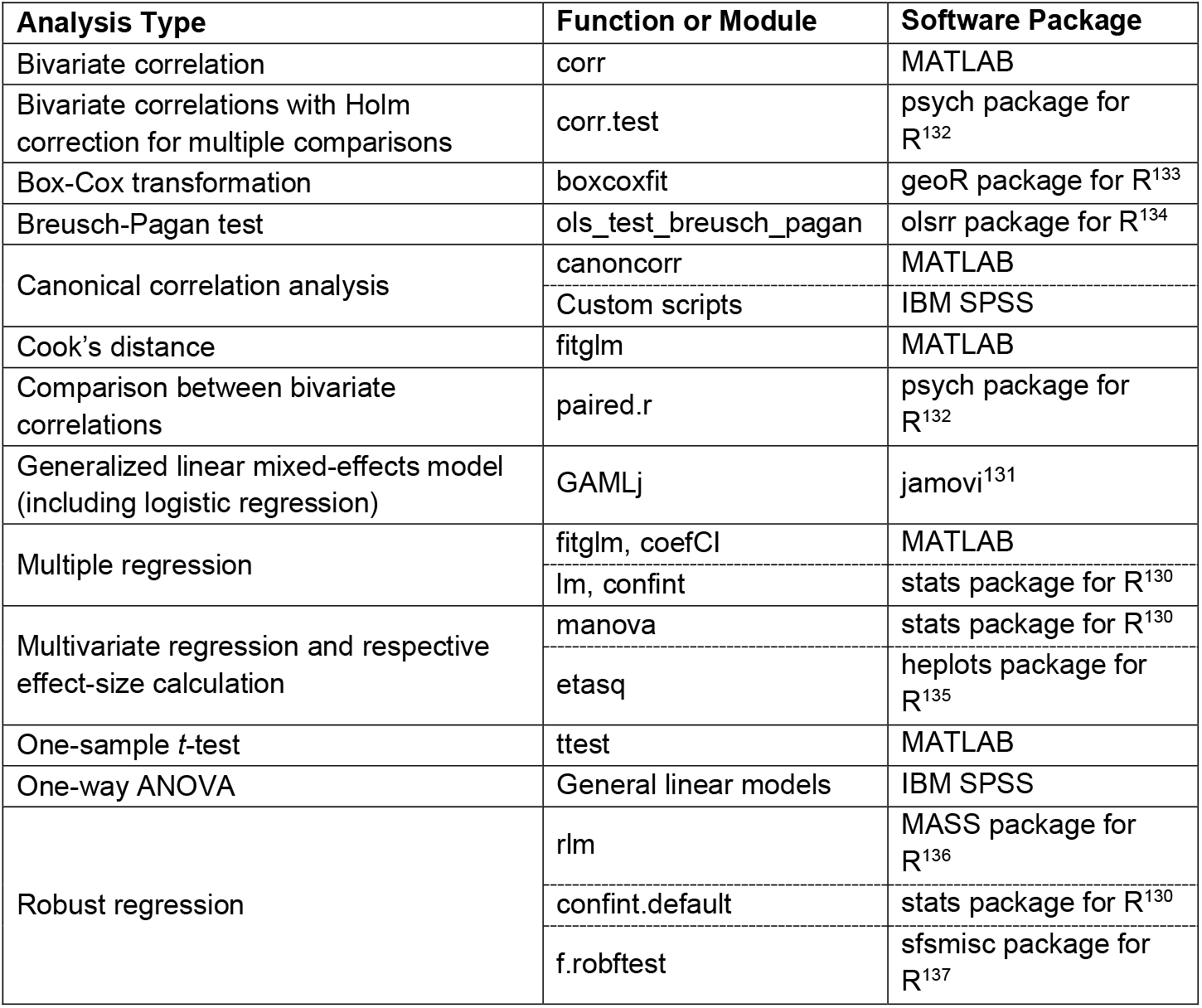
Functions or modules used to conduct each type of statistical analysis. The analysis types are listed alphabetically.

## ACKNOWLEDGEMENTS

We are very grateful to Klaas Stephan for his generous guidance and support of this project. We gratefully acknowledge support by the René and Susanne Braginsky Foundation, the University of Zurich (UZH), the Clinical Research Priority Program “Molecular Imaging” at UZH, Fundação para a Ciência e a Tecnologia, Portugal (PhD fellowship PD/BD/105852/2014 to VAC), the Tourette Association of America (TVM), and the *Brainstorm* Program at the Robert J. & Nancy D. Carney Institute for Brain Science (FHP). VAC and TVM also acknowledge their participation in a twinning project (SynaNet) from the European Union Horizon 2020 Programme (project number: 692340). We also acknowledge the valued study support of our colleagues Thomas Baumgartner, Karl Treiber, Cornelia Schnyder, Andreea Diaconescu, Sandra Iglesias, Helene Haker, and Quentin Huys. Finally, we thank Anderson M. Winkler (University of Texas Rio Grande Valley) for guidance concerning the permutation tests used to obtain *p*-values for the cross-loadings in the canonical correlation analysis.

## EXTENDED DATA FIGURES

**Extended Data Figure 1.**
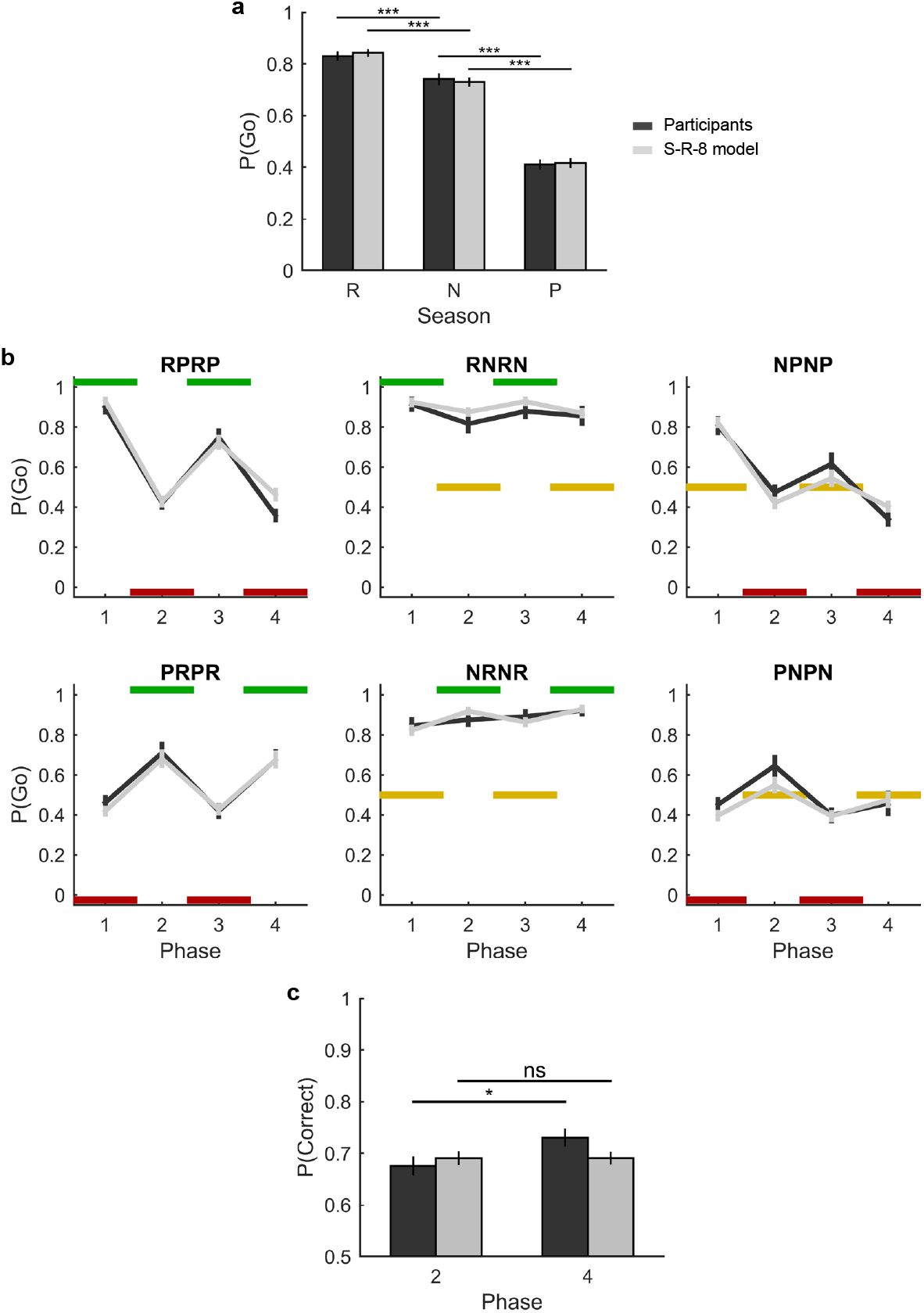
Model S-R-8, the S-R equivalent of winning model S-S-R-8, replicates some of the results concerning participants’ group-level behavior (panels a and b) but not the result that requires state inference (panel c). This figure is equivalent to Fig. 5 in the article but presents the fits of model S-R-8 rather than model S-S-R-8. The average of participants’ behavior is shown in dark gray; the average of the fits of model S-R-8 for individual participants is shown in light gray. **a**, Average (±SEM) of the probability of Go responses, P(Go), exhibited by participants (left bar of each pair; dark gray) and model fits (right bar of each pair; light gray) for each season type: rewarding (R), neutral (N), and punishing (P). We analyzed the model fits using the same approach that we described in the caption of Fig. 5a. Like for participants (see caption of Fig. 5a), P(Go) for the fits of model S-R-8 was largest for rewarding, intermediate for neutral, and smallest for punishing seasons (main effect of season type: 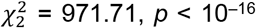; repeated contrasts: rewarding vs. neutral, *b* = 0.68, *z* = 8.79, *p* < 10^−16^, 95% CI [0.53, 0.83], *OR* = 1.97; neutral vs. punishing, *b* = 1.39, *z* = 23.41, *p* < 10^−16^, 95% CI [1.27, 1.50], *OR* = 4.00). **b**, Average (±SEM) P(Go) as a function of the shell phases for each of the six shells, for participants (dark gray) and model fits (light gray). The fits from model S-R-8 track the changes in participants’ choices as the seasons changed. **c**, Average (± SEM) probability of a correct response, P(Correct), in the phases following the first and second identical shell-state transitions—phases 2 and 4, respectively—for participants (dark gray) and model fits (light gray). As in Fig. 5c, P(Correct) was calculated using only R and P seasons, as there was no correct response for N seasons. Participants and model S-S-R-8 exhibited better performance in phase 4 than in phase 2 (see Fig. 5c and its caption), showing that they adapted faster when a state recurred. In contrast, as expected, model S-R-8 did not show a difference in performance between phases 2 and 4 (mean difference = 0.00, *t*_43_ = 0.01, *p* = .994, 95% CI [–0.02, 0.02], Cohen’s *d* = 0.00). Model S-R-8 does not include state inference, so it must learn everything anew whenever the state changes; for this reason, model S-R-8 is unable to adapt faster when a state recurs. ns: not significant; \*\**p* < .01; \*\*\**p* < .001.

**Extended Data Figure 2.**
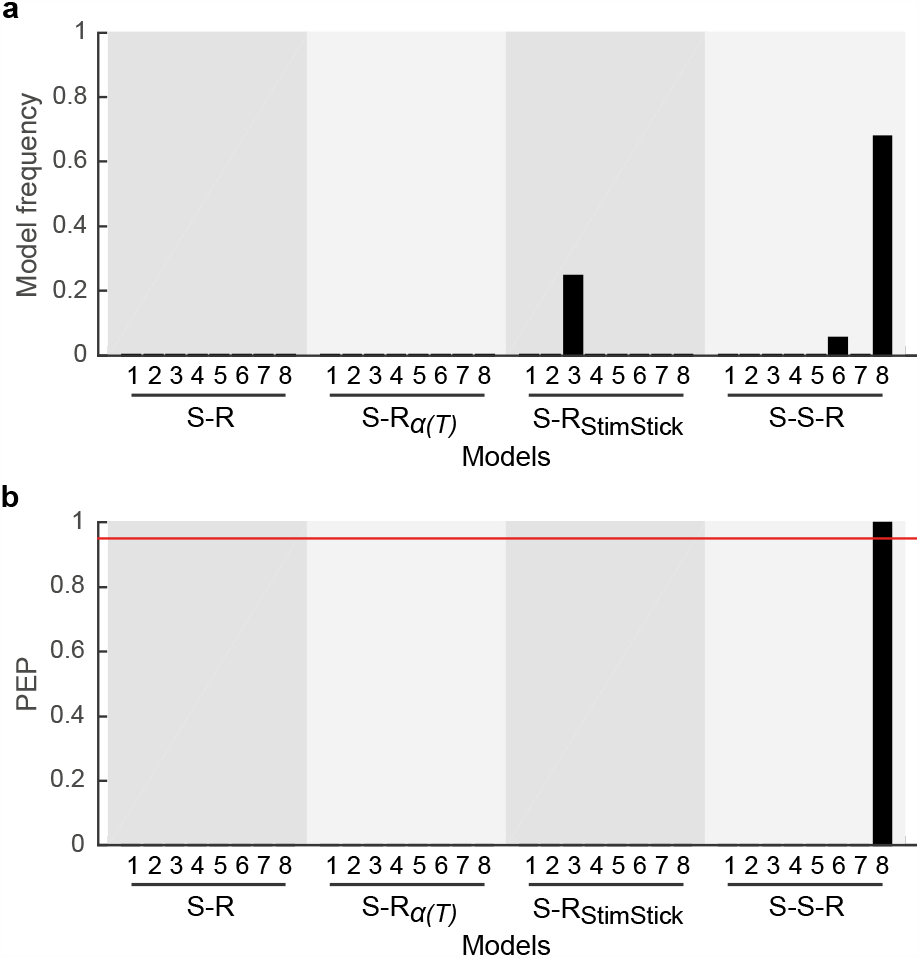
Behavior was explained better by a model that extended stimulus-response learning with state inference (model S-S-R-8) than by models that implemented learning rates that could vary across the task (S-R_α(*T*)_ models) or models that implemented stimulus stickiness (S-R_StimStick_ models; see *Methods – Computational models – Alternative model families*). **a**, Model frequencies. **b**, Protected exceedance probabilities (PEPs). The red line indicates the threshold for confident selection of a model. State-inference model S-S-R-8 was confidently selected as the best model.

