## Supplementary results for "Serotonin Reduces Belief Stickiness"

Vasco A. Conceição*, Frederike H. Petzschner*, David M. Cole, Katharina V. Wellstein, Daniel Müller, Sudhir Raman, Tiago V. Maia

*Authors contributed equally

### EFFECTS OF $\gamma$ AND $\zeta$ ON TASK PERFORMANCE

#### Effects of $\gamma$ on task performance

There is an inherent tradeoff between flexibility and stability of beliefs about a shell's hidden states, which our formulation of belief pliability in terms of $\gamma$ [equation (3) in the *Methods*] adequately captures. On one extreme, maximal, excessive flexibility—better characterized as belief instability—would correspond to always ignoring the previous evidence concerning the state that one is in, determining the distribution of beliefs about the current state, $p_t(h_i)$, using only the current trial's reinforcement, $r_t$. In our formulation, this extreme would correspond to $\gamma$ = 0: this value would cause $p_t'(h_i)$ to become a uniform distribution on every trial [equation (3) in the *Methods*], thereby leading the posterior belief, $p_t(h_i)$, to be fully determined by $r_t$ [equation (4) in the *Methods*]. This extreme would ignore the temporal stability of seasons, which do not change on every trial, causing excessive belief switching. On the other extreme, maximal, excessive belief stickiness corresponds to treating the environment as stationary, disregarding the possibility that the shell's season may change. On this extreme, the current trial's reinforcement, $r_t$, is accorded as much weight as past reinforcements in determining the current beliefs about the season's hidden state. In our formulation, this extreme would correspond to $\gamma$ = 1: this value would cause $p_t'(h_i)$ to equal the previous beliefs, $p_{t-1}(h_i)$ [equation (3) in the *Methods*], thereby leading the posterior belief, $p_t(h_i)$, to incorporate $r_t$ as just another piece of information, accorded as much weight as older information [equation (4) in the *Methods*]. This extreme would ignore the fact that seasons do occasionally change—as participants are explicitly told in the task instructions—thereby causing excessive belief stickiness. (An even more extreme case of belief stickiness is obtained using $\zeta$, as described in the next section.)

From the foregoing, it should be clear that $\gamma$ should have a nonmonotonic effect on task performance. To better characterize this effect, we varied $\gamma$ and assessed the effect on task performance in 1,000 simulations. We characterized task performance as the number of correct responses (Go responses in rewarding seasons and NoGo responses in punishing seasons). In each of the 1,000 simulations, we drew a random sample of $\gamma$ from a uniform distribution between 0 and 1 (the range of $\gamma$). With $\gamma$ set to the sampled value, we then had the model perform each of the 44 traces of the task that were provided to the 44 participants. We always set the value of each of the other parameters to its corresponding mean value across participants. Finally, we averaged the number of correct responses in the 44 traces; this average produced a datapoint associated with that simulation. In this way, we obtained 1,000 datapoints (Supplementary Results Fig. 1a).

We analyzed the datapoints thus obtained using polynomial regression (centering $\gamma$). We tested cubic, quadratic, and linear models. In each model, we also included all lower-order polynomial terms (e.g., in the cubic model, we included cubic, quadratic, and linear terms, in addition to the intercept). All models were vastly superior to the null (i.e., intercept-only) model (cubic model: $F_{4,996}$ = 626.23, $p < 10^{-227}$; quadratic model: $F_{3,997}$ = 816.42, $p < 10^{-209}$; linear model: $F_{2,\,998}$ = 1,543.60, $p < 10^{-204}$). In the cubic model, the coefficients for cubic, quadratic, and linear terms were all highly significant (cubic: $b$ = 15.48, $t_{996}$ = 9.69, $p < 10^{-20}$, 95% CI [12.34, 18.61]; quadratic: $b$ = –2.28, $t_{996}$ = –5.49, $p < 10^{-7,}$ 95% CI [–3.10, –1.47]; linear: $b$ = –6.91, $t_{996}$ = –25.86, $p < 10^{-112,}$ 95% CI [–7.43, –6.38]). In addition, the cubic model had a larger adjusted $R^2$ and smaller Akaike Information Criterion (AIC) and Bayesian Information Criterion (BIC) than both the quadratic and linear models (Supplementary Results Table 1, top). We therefore retained the cubic model.

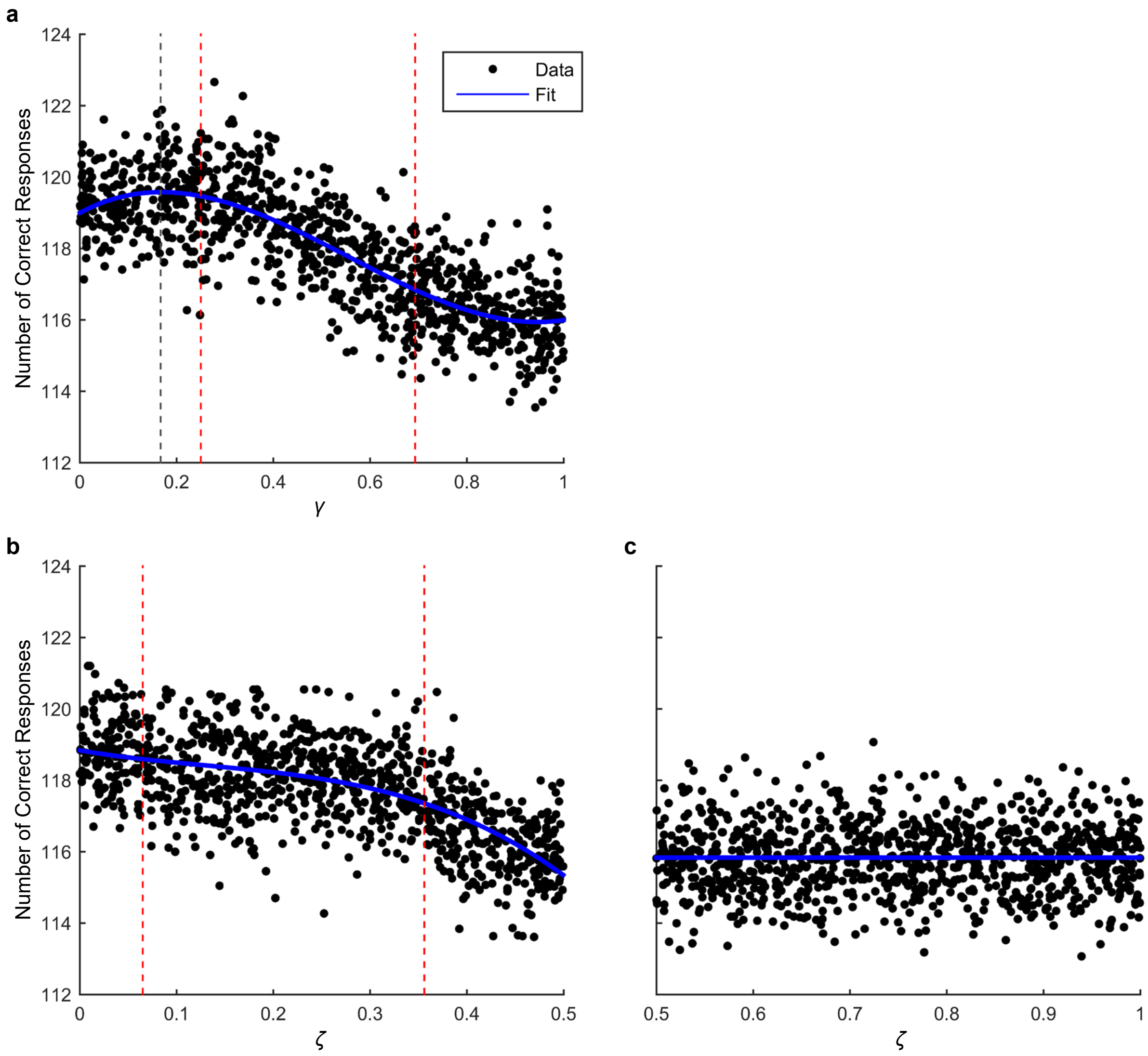

**Supplementary Results Fig. 1** Results of the simulations assessing the effects of varying $\gamma$ and $\zeta$ on the overall number of correct responses in the task. **a,** Results for $\gamma$. Each datapoint corresponds to one simulation. The blue line shows the fit using cubic polynomial regression (which was better than quadratic polynomial regression, linear regression, or an intercept-only model; see text). The dashed vertical gray line shows the value of $\gamma$ that produces the largest number of correct responses (corresponding to a zero of the derivative of the cubic fit). The left and right dashed vertical red lines indicate the minimum and maximum values of $\gamma$, respectively, for participants in our sample. The portion of the fit curve (blue line) between the dashed red lines decreases monotonically, showing that, for the values of $\gamma$ obtained for participants in our sample, larger values of $\gamma$ were detrimental to performance—i.e., corresponded to detrimental belief stickiness. **b,** Results for $\zeta$, sampled between 0 and 0.5. The blue line shows the fit using cubic polynomial regression (which was better than quadratic polynomial regression, linear regression, or an intercept-only model; see text). The left and right dashed vertical red lines indicate the minimum and maximum values of $\zeta$, respectively, for participants in our sample. The number of correct responses decreases monotonically with increasing values of $\zeta$ in the entire range from 0 to 0.5. **c,** Results for $\zeta$, sampled between 0.5 and 1. The flat horizontal blue line shows the fit using an intercept-only model (which was better than cubic polynomial regression, quadratic polynomial regression, or linear regression; see text). Values of $\zeta$ between 0.5 and 1 all lead to the same impairment in task performance.

| **Model** | **Adjusted $R^2$** | **AIC** | **BIC** |
|---|---|---|---|
| Simulations of $\boldsymbol{\gamma}$ $(\mathbf{0} < \boldsymbol{\gamma} < \mathbf{1})$ | | | |
| Cubic | 0.652 | 2,834 | 2,854 |
| Quadratic | 0.620 | 2,922 | 2,937 |
| Linear | 0.607 | 2,955 | 2,965 |
| Simulations of $\boldsymbol{\zeta}$ $(\mathbf{0} < \boldsymbol{\zeta} < \mathbf{0.5})$ | | | |
| Cubic | 0.451 | 2,891 | 2,911 |
| Quadratic | 0.446 | 2,899 | 2,914 |
| Linear | 0.402 | 2,975 | 2,984 |

**Supplementary Results Table 1** Model fits for the cubic, quadratic, and linear regressions of the simulations of task performance as a function of $\gamma$ (top) and $\zeta$ (bottom), with $\gamma$ ranging between 0 and 1, and with $\zeta$ ranging between 0 and 0.5. Larger values of adjusted $R^2$ but smaller values of the Akaike Information Criterion (AIC) and Bayesian Information Criterion (BIC) indicate better fits (while penalizing for model complexity).

As expected, for small values of $\gamma$, the number of correct responses increased as $\gamma$ increased, but then, as $\gamma$ continued to increase, the number of correct responses steadily declined (Supplementary Results Fig. 1a). At first sight, this nonmonotonic effect of $\gamma$ on task performance might seem to question our (simplified) characterization of $\gamma$, in the main text and Methods, as a detrimental belief-stickiness parameter. However, all values of $\gamma$ for the participants in our sample were in the monotonically decreasing portion of the curve (Supplementary Results Fig. 1a). In other words, for the values of $\gamma$ observed in our sample, larger values of $\gamma$ did always correspond to excessive belief stickiness that hampered task performance. We therefore chose to characterize $\gamma$ as a detrimental belief-stickiness parameter in the main text and Methods to make exposition more accessible to readers who might be less interested in all the technical nuances. This characterization, while somewhat simplified if one considers all possible values of $\gamma$ in the allowed range ($0 < \gamma < 1$), was entirely accurate for the values observed in our dataset.

For the same reason, we need not concern ourselves with the nonmonotonic effect of $\gamma$ on task performance, in the full allowed range of $\gamma$ (from 0 to 1), when analyzing the relation of $\gamma$ to other variables in our sample (such as escitalopram level or group or OCI-R scores). In the observed range of values of $\gamma$, the effects of $\gamma$ on task performance were monotonic.

**Effects of $\zeta$ on task performance**

The parameter $\zeta$ supports a slightly different, arguably even more extreme, form of belief stickiness than that afforded by high values of $\gamma$ alone. As noted in the previous section, a value of $\gamma$ close to 1 (i.e., to the maximum value) is equivalent to assuming a stationary environment. Even with such an extreme value of $\gamma$, however, incoming information (the observed reinforcement, $r_t$, on each trial) would still be incorporated into the posterior beliefs about the hidden states, $p_t(h_i)$ [equation (4) in the *Methods*]: it is only that new information would be afforded the same weight as previous, possibly outdated information. The parameter $\zeta$, however, is added directly to the inference about the current state, causing a tendency to believe that the state remains the same as it was previously, even when the posterior beliefs, $p_t(h_i)$, indicate otherwise [equation (5) in the *Methods*]. This tendency therefore ignores incoming information (and it is in that sense that we say that it can be an even more extreme form of belief stickiness).

From the foregoing, it is intuitive that larger values of $\zeta$ should be more detrimental to task performance. To test this intuition, we varied $\zeta$ and assessed the effect on the number of correct responses through simulations, like we did for $\gamma$. The simulations were entirely parallel to the ones for $\gamma$, but varying $\zeta$ instead of $\gamma$.

In a first set of 1,000 simulations, we sampled $\zeta$ from a uniform distribution between 0 and 0.5 (the range we set for $\zeta$). Like we did for $\gamma$, we analyzed the datapoints thus obtained (Supplementary Results Fig. 1b) using cubic, quadratic, and linear models (centering $\zeta$ and including, in each model, all lower-order polynomial terms). All models were vastly superior to the null (i.e., intercept-only) model (cubic model: $F_{4,\,996} = 274.67$, $p < 10^{-129}$; quadratic model: $F_{3,997} = 403.58$, $p < 10^{-128}$; linear model: $F_{2,\,998} = 672.97$, $p < 10^{-113}$). In the cubic model, the coefficients for cubic, quadratic, and linear terms were all highly significant (cubic: $b = -42.06$, $t_{996} = -3.12$, $p = .002$, 95% CI [–68.50, –15.63]; quadratic: $b = -15.11$, $t_{996} = -8.90$, $p < 10^{-17,}$ 95% CI [–18.44, –11.78]; linear: $b = -4.35$, $t_{996} = -7.72$, $p < 10^{-13,}$ 95% CI [–5.45, –3.24]). In addition, the cubic model had a larger adjusted $R^2$

and smaller AIC and BIC than both the quadratic and linear models (Supplementary Results Table 1, bottom). We therefore retained the cubic model. The number of correct responses decreased monotonically with increasing values of $\zeta$ in the entire range from 0 to 0.5.

A reasonable question based on our mathematical formulation of the effect of $\zeta$ [equation (5) in the *Methods*] is why we restricted $\zeta$ to be between 0 and 0.5 rather than between 0 and 1. After all, only a value of $\zeta$ = 1 would seem to ensure complete belief stickiness—i.e., that the model would always believe that the hidden state remained the same, regardless of any incoming information. In our task, however, a value of $\zeta$ equal to 0.5 is sufficient to, in practice, virtually always prevent changes in the belief about the current state [because the difference in posterior beliefs, $p_t(h_i)$, between the belief for the previous hidden state and the belief for the current maximum *a posteriori* (MAP) state is virtually always less than 0.5]. Allowing the range of $\zeta$ to extend upwards of 0.5, until 1, would therefore cause problems with model identifiability, as a large range of values of $\zeta$ would lead to indistinguishable behavior. To demonstrate this effect, we conducted another set of 1,000 simulations varying $\zeta$ and assessing the effect on the number of correct responses, but sampling $\zeta$ from a uniform distribution between 0.5 and 1 (Supplementary Results Fig. 1c).

Like we did for $\gamma$ and for the simulations of $\zeta$ in the range of 0 to 0.5, we fit cubic, quadratic, and linear models to the resulting datapoints (centering $\zeta$ and including, in each model, all lower-order polynomial terms). None of these models were significantly better than the null, intercept-only model (cubic model: $F_{4,\ 996}$ = 0.43, $p$ = .735; quadratic model: $F_{3,997}$ = 0.63, $p$ = .532; linear model: $F_{2,\ 998}$ = 0.36, $p$ = .551). Moreover, the adjusted $R^2$ for all these models was very close to 0 and even negative (cubic model: adjusted $R^2$ = –0.002; quadratic model: adjusted $R^2$ = –0.001; linear model: adjusted $R^2$ = –0.001). These results are consistent with the idea that values of $\zeta$ between 0.5 and 1 all lead to the same behavior, as we had hypothesized. Indeed, the resulting datapoints seem to fall exactly around a horizontal flat line corresponding to their mean value (i.e., the intercept-only model; Supplementary Results Fig. 1c). This set of simulations therefore confirms that values of $\zeta$ equal to 0.5 already produce maximal belief stickiness in our task; allowing values of $\zeta$ to range up to 1 would therefore cause identifiability problems.

### MODEL- AND PARAMETER-RECOVERY SIMULATIONS

#### Model recovery

We tested whether we could successfully recover our candidate models using our task, sample size, and model-inversion and model-comparison procedures. For this purpose, we generated synthetic data with different models and tested whether, by following our procedures, we would accurately recover which model had generated the data. Given that the model-inversion process was extremely intensive computationally and that the computation time was proportional to the square of the number of models, we used for these simulations a subset of 8 of the 16 candidate models (S-R-1, S-R-4, S-R-5, S-R-8, S-S-R-1, S-S-R-4, S-S-R-5, and S-S-R-8). For each of these 8 models, we generated a dataset of synthetic data for 44 simulated participants, using the 44 task traces presented to the real participants (and a fixed set of parameters; Supplementary Results Table 2). For each such dataset, we applied our model-inversion and model-comparison procedures to determine if we could recover the model used to generate the data with a protected exceedance probability (PEP) > 0.95—the same criterion that we used for the actual data[1]. We found that we successfully recovered the model used to generate the data with 100% accuracy (Supplementary Results Fig. 2a). We obtained similar results using other parameter sets (not shown).

| **Model** | $\boldsymbol{\alpha^{+}}$ | $\boldsymbol{\alpha^{-}}$ | $\boldsymbol{\beta^{+}}$ | $\boldsymbol{\beta^{-}}$ | $\boldsymbol{\varphi}$ | $\boldsymbol{\gamma}$ | $\boldsymbol{\zeta}$ |
|---|---|---|---|---|---|---|---|
| S-R-1 | 0.2 | | 2 | | | | |
| S-R-4 | 0.3 | 0.1 | 3 | 1 | | | |
| S-R-5 | 0.5 | | 2 | | 0.2 | | |
| S-R-8 | 0.3 | 0.1 | 3 | 1 | 0.2 | | |
| S-S-R-1 | 0.2 | | 2 | | | 0.5 | 0.1 |
| S-S-R-4 | 0.3 | 0.1 | 3 | 1 | | 0.5 | 0.1 |
| S-S-R-5 | 0.2 | | 2 | | 0.2 | 0.5 | 0.1 |
| S-S-R-8 | 0.3 | 0.1 | 3 | 1 | 0.2 | 0.5 | 0.1 |

**Supplementary Results Table 2** Parameters used in the model-recovery simulations for each of the simulated models. Note that models S-R-1 and S-R-5, and therefore also models S-S-R-1 and S-S-R-5, have a single value for $\alpha$ and a single value for $\beta$ (Methods Table 1). Not all parameters are present in all models (Methods Table 1); we shaded in gray those that are not. For each parameter, we always used the same value in all models that included that parameter.

### Parameter recovery

We ran an additional set of simulations to test whether we could also accurately recover the model parameters. For these simulations, we used only the more complex model, S-S-R-8, because all other models are special cases of this model (*Methods – Computational models*). We generated 7 datasets, each of which varied 1 of the 7 parameters while keeping the other 6 parameters fixed. For each dataset, we created 44 simulated participants, using the 44 task traces presented to the real participants. In each dataset, each simulated participant had a different value of the parameter being varied (with the specific value sampled randomly from the prior distribution for that parameter), but all simulated participants had the same values for the remaining parameters (set to the fitted group mean: $\alpha^{+}$ = 0.411*;* $\alpha^{-}$ = 0.559; $\beta^{+}$ = 3.815; $\beta^{-}$ = 3.516; $\varphi$ = 0.504; $\gamma$ =

0.456; $\zeta$ = 0.210). We then inverted the model, with all parameters free to vary. This procedure allowed us to address two questions. First, we assessed whether we could recover individual parameters accurately, by measuring, for each dataset, the correlation between the recovered values of the parameter that was varied and the parameter values used during simulation. Second, we assessed whether variation in a parameter would be erroneously picked up in other parameters during model inversion, by assessing whether recovered parameters that had not been varied in a dataset correlated with the value of the parameter that had been varied in that dataset. We were particularly interested in the belief-stickiness parameters, as those were the main focus of our analyses.

All parameters could be recovered accurately (Supplementary Results Fig. 2b). Moreover, the strongest correlation was always between the varied parameter and the estimated values of the same parameter (Supplementary Results Fig. 2c–d), showing that parameter variation was always captured most strongly in the correct parameter. At least in this set of simulations, with the parameter values used, there was some crosstalk between S-R parameters, with variation in several S-R parameters being erroneously picked up in other S-R parameters (Supplementary Results Fig. 2c). The crosstalk obtained in these simulations might conceivably be reduced with additional simulations using different values for the fixed parameters; still, such crosstalk is common in reinforcement-learning models[2,3]. Crucially for our purposes, however, crosstalk between S-R parameters and the belief-stickiness parameters was limited (Supplementary Results Fig. 2c–d). Our design and models therefore largely orthogonalized the S-R parameters from the belief-stickiness parameters. More importantly, we can be confident that variation in the fitted belief-stickiness parameters likely reflects true variation in belief stickiness (especially if such variation is unaccompanied by variation in S-R parameters, as was the case in our findings).

The two belief-stickiness parameters were not clearly distinguishable from each other (Supplementary Results Fig. 2c–d). As noted in the main text, however, we designed these two parameters to tap into the same underlying construct of belief stickiness, so we were not concerned with distinguishing between them. Moreover, in all analyses of these two parameters, we used techniques that considered the parameters' joint effects.

### REPLICATION OF THE POSITIVE ASSOCIATION BETWEEN ESCITALOPRAM PLASMA LEVEL AND STATE INFERENCE

We replicated the positive association between escitalopram plasma level and adequate state inference with two approaches that did not use computational models. Although these analyses are cruder than those that used computational models, they provide convergent evidence for the positive association between escitalopram plasma level and state inference.

**a**

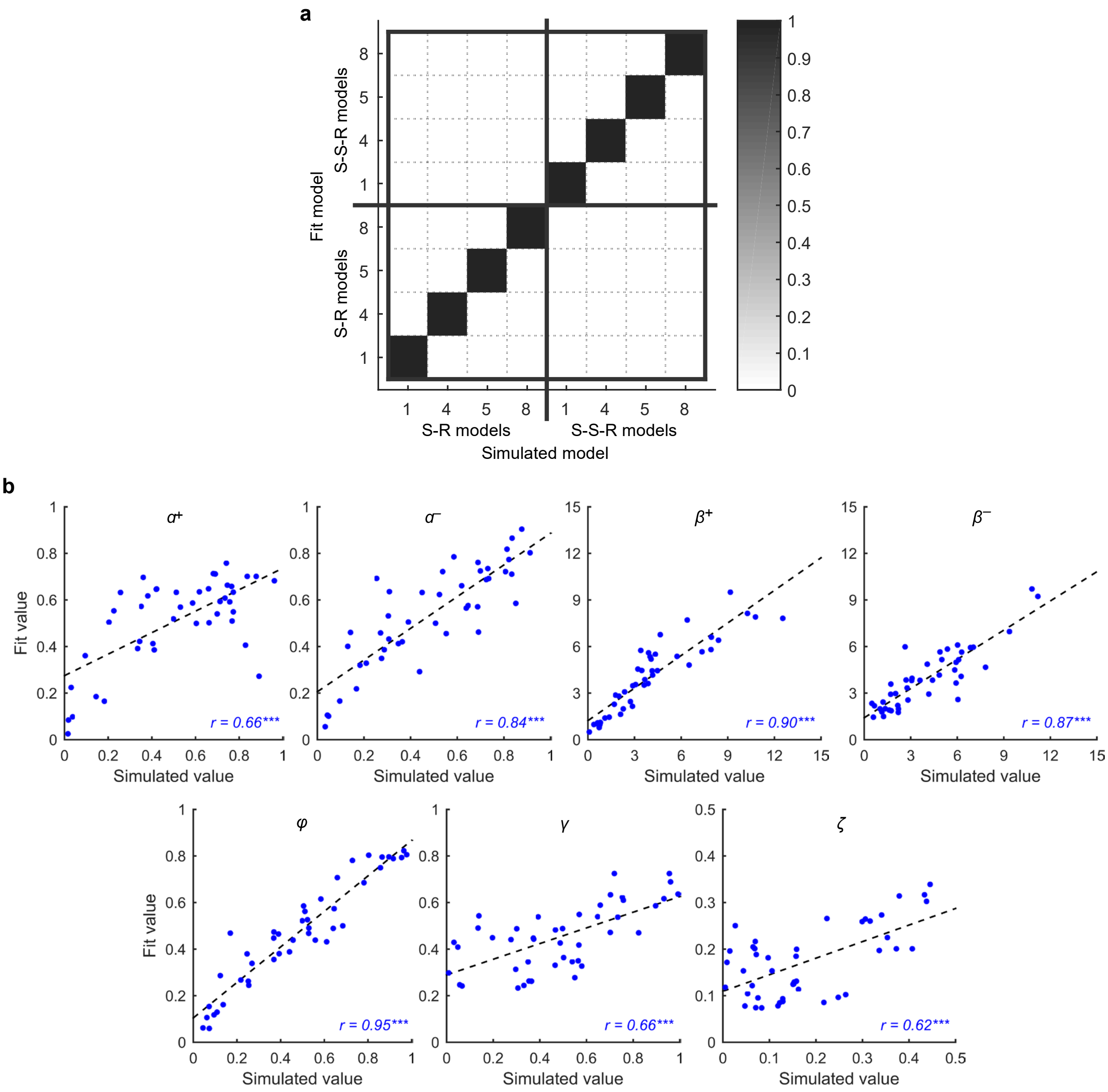

**c**

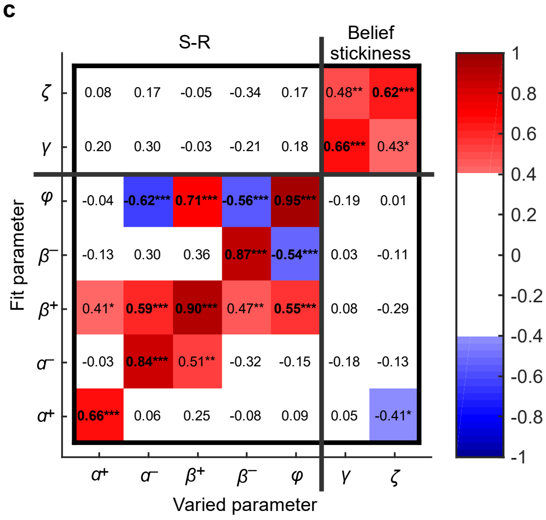

**d**

| Varied parameter | $t_{41}$ | $p$ |
|---|---|---|
| $\zeta$ | 2.16 | .037 |
| $\gamma$ | 1.97 | .055 |
| $\varphi$ | 6.28 | $< 10^{-6}$ |
| $\beta^-$ | 3.63 | .001 |
| $\beta^+$ | 3.30 | .002 |
| $\alpha^-$ | 2.58 | .013 |
| $\alpha^+$ | 2.03 | .049 |

**Supplementary Results Fig. 2** Model- and parameter-recovery results. **a,** Confusion matrix[4] for the model-recovery simulations. The matrix shows, for each model that was simulated (corresponding to one column), the protected exceedance probabilities (PEPs) for the 8 models that were fitted (each of which is represented in one row). The fact that the matrix is diagonal shows that all simulated models could be recovered with 100% accuracy. **b,** Scatterplots showing the relation between the simulated values of each parameter and the fitted values of the same parameter. Each plot corresponds to one of the 7 datasets, in which only the indicated parameter was varied while the other parameters were fixed. The dotted lines represent linear fits. Pearson correlations (*r*) are shown at the bottom of each plot. Parameters could be accurately recovered: the correlation between the simulated and fit values was large and highly significant ($r_{42} > 0.65$, $p < .001$) in each of the 7 datasets. **c,** Pearson correlations between the simulated values of the parameter that was varied in each dataset and the fitted values of all parameters. Each column corresponds to one of the 7 datasets, in which only the parameter indicated in the column label was varied while the other parameters were fixed. The plots shown in panel b correspond to the diagonal of this matrix. Significant correlations are color-coded according to the value of *r*, as shown in the bar on the right; nonsignificant correlations are shown in white. Within each dataset, we corrected for multiple comparisons using the Holm procedure. For each dataset, the correlation was always strongest between the varied parameter and itself: the largest absolute value of the correlations in each column falls on the diagonal. Crosstalk between S-R and belief-stickiness parameters was limited, as evidenced by the fact that the two top rows and the two right columns are nearly always white outside of the 2 x 2 square at the top right. **d,** Statistical comparison, for each varied parameter, of the correlation of that parameter with itself versus the next strongest correlation (i.e., the next correlation with the largest absolute value) for that parameter. In other words, for each column of the matrix in panel c, we compared the correlation in the diagonal with the next largest correlation (in absolute value) in that column. For all parameters except $\gamma$, the correlation of the parameter with itself was significantly greater than the next largest correlation (in absolute value) for that parameter. For $\gamma$, the correlation between $\gamma$ and itself was only marginally significantly greater than the correlation between $\gamma$ and $\zeta$, consistent with the finding that $\gamma$ and $\zeta$ were not clearly distinguishable (panel c). This difficulty distinguishing $\gamma$ and $\zeta$ was expected because both parameters were designed to tap into the same underlying construct of belief stickiness. The correlation between $\gamma$ and itself was, however, significantly greater than the largest (absolute) correlation between $\gamma$ and one of the S-R parameters ($\varphi$), $t_{41} = 2.88$, $p = .006$, consistent with the idea that the belief-stickiness parameters ($\gamma$ and $\zeta$) could be distinguished from the S-R parameters (panel c). *$p < .05$; **$p < .01$; ***$p < .001$.

### Escitalopram plasma levels related positively to the increase in performance from phase 2 to phase 4

As noted in the main text, adequate state inference fosters an increase in performance—i.e., in the probability of a correct response—from phase 2 to phase 4 (Fig. 1d). Phases 2 and 4 are directly comparable because the season is the same in both, and the preceding season is also the same in both. If participants adequately infer the shell states, however, in phase 4 they can reuse the knowledge that they acquired in phase 2, thereby adapting behavior more quickly. The increase in the probability of a correct response from phase 2 to phase 4 can therefore serve as a (somewhat crude) proxy for adequate state inference. To investigate the relation of this state-inference proxy with escitalopram (level or group) and OCI-R scores (OCI-R obsessing and OCI-R other), we regressed the increase in the probability of a correct response from phase 2 to phase 4 on group, escitalopram plasma level, OCI-R obsessing, and OCI-R other.

The coefficient for the escitalopram plasma level was positive and strongly significant ($b$ = 0.006, $t_{38}$ = 3.07, $p$ = .004, 95% CI [0.002, 0.009]; Supplementary Results Fig. 3a). This finding replicates the positive association between escitalopram plasma level and state inference, using the increase in the probability of a correct response from phase 2 to phase 4 as a proxy for state inference.

Neither the coefficient for OCI-R obsessing nor the coefficient for OCI-R other were significant ($b$ = –0.003, $t_{38}$ = –0.201, $p$ = .842, 95% CI [–0.030, 0.025], and $b$ = –0.001, $t_{38}$ = –0.250, $p$ = .804, 95% CI [–0.009, 0.007], respectively). Although OCI-R obsessing and OCI-R other correlated strongly and significantly ($r_{41}$ = 0.69, $p < 10^{-6}$), their variance inflation factors were below 5 (VIF = 1.93 and VIF = 1.90, respectively), suggesting that the lack of significance of each was not due to its correlation with the other. Indeed, with backward elimination of OCI-R other, OCI-R obsessing remained nonsignificant ($b$ = –0.005, $t_{39}$ = –0.52, $p$ = .608, 95% CI [–0.025, 0.015]), and with backward elimination of OCI-R obsessing, OCI-R other remained nonsignificant ($b$ = –0.002, $t_{39}$ = –0.54, $p$ = .593, 95% CI [–0.007, 0.004]). Thus, OCI-R scores, including OCI-R obsessing, did not relate to the increase in the probability of a correct response from phase 2 to phase 4. This null finding likely reflects the crudeness of this behavioral measure as a proxy for state inference, as in the more accurate and sensitive analyses based on computational models, we found a highly significant negative relation between OCI-R obsessing and state inference ($p$ = .002).

The coefficient for group was also not significant ($b$ = 0.010, $t_{38}$ = 0.24, $p$ = .814, 95% CI [–0.073, 0.092]). Although this finding might seem surprising considering the significant effect of escitalopram level, it can be explained if only sufficiently high escitalopram levels facilitate state inference. As noted above, in these analyses, the coefficient for group represents the difference between the escitalopram group, at its mean level of escitalopram, and the placebo group. If only escitalopram levels that tended to be higher than the mean levels facilitated state inference, we would not expect a significant effect for the group coefficient. This null finding for group mimics the null finding for group in the analyses using computational models, which we address in detail in the main text.

One possible concern with the preceding regression is whether the significant effect of escitalopram level is unduly influenced by the subject with particularly high escitalopram level, who also had a particularly large increase in the probability of a correct response from phase 2 to phase 4 (see the rightmost and uppermost data point in Supplementary Results Fig. 3a). We therefore checked whether this datapoint was unduly influential; it was not (Cook's distance $D$ = 0.22, below the commonly recommended absolute cutoffs of 1 or 0.5 to identify influential points[5]). Still, out of an abundance of caution, we also replicated this analysis using robust regression (with bisquare weighting). The robust regression produced the same conclusions in terms of statistical significance, with the coefficient for escitalopram level even becoming slightly more significant ($b$ = 0.006, $t_{38}$ = 3.15, $p$ = .003, 95% CI [0.002, 0.010]).

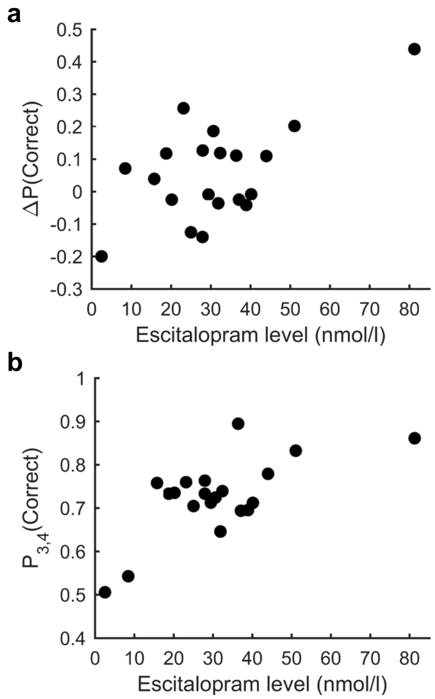

**Supplementary Results Fig. 3** Replication of the positive association between escitalopram plasma level and state inference using simpler measures that do not include computational models. **a,** Relation between escitalopram plasma levels and the change in the probability of a correct response from phase 2 to phase 4, ΔP(Correct), which is a behavioral proxy for adequate state inference. ΔP(Correct) was calculated using only R and P seasons because is no correct response for N seasons. Higher escitalopram levels were associated with larger values of ΔP(Correct) and therefore with better state inference. **b,** Relation between escitalopram plasma levels and the probability of a correct response in phases 3 and 4, $P_{3,4}$(Correct), which involve state revisiting. Like ΔP(Correct), $P_{3,4}$(Correct) was calculated using only R and P seasons because there is no correct response for N seasons. Higher escitalopram levels were associated with a greater probability of a correct response in phases 3 and 4.

### Escitalopram plasma level related positively to performance in later phases, which revisit states

Better state inference should improve performance—i.e., the probability of a correct response—in phases 3 and 4, which revisit states and therefore benefit especially from better state inference, to support reutilization of the previously learned contingencies. To test whether performance in phases 3 and 4 related to escitalopram (level or group) and OCI-R scores, we regressed the probability of a correct response in phases 3 and 4 on group, plasma escitalopram level, OCI-R obsessing, and OCI-R other.

The coefficient for the plasma escitalopram level was positive and strongly significant ($b$ = 0.004, $t_{38}$ = 3.39, $p$ = .002, 95% CI [0.002, 0.006]; Supplementary Results Fig. 3b), showing that increased escitalopram levels were associated with a greater probability of a correct response in phases 3 and 4. Neither the coefficient for OCI-R obsessing nor the coefficient for OCI-R other were significant ($b$ = –0.001, $t_{38}$ = –0.09, $p$ = .931, 95% CI [–0.017, 0.016], and $b$ = –0.001, $t_{38}$ = –0.62, $p$ = .539, 95% CI [–0.006, 0.003], respectively). The coefficient for group was also not significant ($b$ = 0.019, $t_{38}$ = 0.80, $p$ = .429, 95% CI [–0.030, 0.069]). As in the regression in the previous section, the subject with particularly high escitalopram level also had a relatively large probability of a correct response in phases 3 and 4 (Supplementary Results Fig. 3b). Again, though, this datapoint was not unduly influential (Cook's distance $D$ = 0.48), and robust regression (with bisquare weighting) produced the same conclusions in terms of statistical significance ($b$ = 0.004, $t_{38}$ = 3.22, $p$ = .003, 95% CI [0.001, 0.006]).

The results in this section mimic those in the previous section. Considered together, these results provide further converging evidence that larger escitalopram plasma levels improved state inference, which resulted in better performance in the phases that involved state revisiting (phases 3 and 4).

Neither the analysis in this section nor that in the previous section found a significant effect of OCI-R obsessing. These non-significant results contrast with those from the model-based analyses in the main text, in which the effect of OCI-R obsessing was highly significant. The impurity of direct behavioral measures, such as performance, reduces their sensitivity and specificity to detect the processes of interest, when compared to analyses that use computational models[6]. The superiority of analyses based on computational models is illustrated, for example, by their ability to predict treatment outcome when analyses based on direct behavioral measures fail[7]. We therefore place substantially more weight on the results of the analyses that used computational models.

### ESCITALOPRAM DID NOT RELATE TO S-R PARAMETERS

#### Escitalopram plasma levels predominantly affected state inference rather than a Go bias

As noted in the main text, earlier work found that citalopram increased the Go bias[8], but our model-based analyses did not detect an effect of escitalopram on the Go bias. To investigate more closely whether escitalopram had such an effect in our task, in addition to, or instead of, an effect on state inference, we assessed how the probability that participants would perform a Go response, P(Go), related to escitalopram (level or group) during the different season types. For this purpose, we used a trial-level mixed-effects logistic regression, with participants' choices (Go or NoGo, coded as 1 or 0, respectively) as the dependent variable, and with group, escitalopram plasma level, season type (rewarding, neutral, punishing), and the interactions of escitalopram plasma level and of group with season type as the independent variables (and with the intercept as a random effect to capture participants' possible different propensities to perform Go). If escitalopram increased the Go bias, as suggested by the prior work, we would expect to see a main effect of escitalopram (level and/or group) but no interaction between escitalopram (level and group) and season type. If instead, or in addition, escitalopram facilitated state inference, we would expect to see an interaction between escitalopram (level or group) and season type, as better state inference affects P(Go) differently in the different season types. In fact, if escitalopram selectively, or at least predominantly, facilitated state inference, its plasma levels should relate with P(Go) positively for the rewarding seasons, and possibly also for neutral seasons (in which shell collection does not yield rewards but provides state information), but negatively for the punishing seasons.

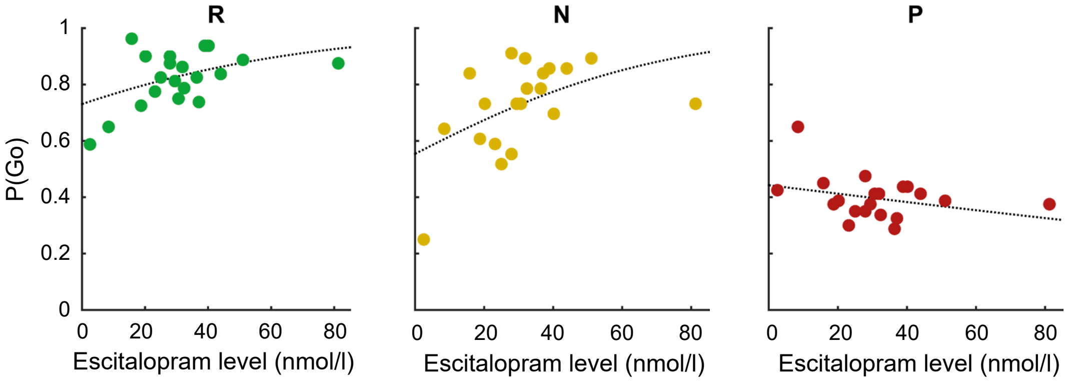

**Supplementary Results Fig. 4** Relation between the probability of a Go response, P(Go), and escitalopram plasma levels for the different season types [from left to right: rewarding (R), neutral (N), and punishing (P)]. The dotted lines represent the fit using a trial-level mixed-effects logistic regression, with participants' choices (Go or NoGo) as the dependent variable and with escitalopram level, season type, and their interaction as the independent variables. Higher escitalopram levels were associated with increased P(Go) in R and N seasons and also, albeit non-significantly, with decreased P(Go) in P seasons, as predicted if higher escitalopram levels facilitated state inference rather than affecting the Go bias. *$p$ < .05. ***$p$ < .001.

Consistent with an effect of escitalopram level on state inference, we found an interaction between season type and escitalopram level ($\chi^2_2$ = 37.15, $p < 10^{-8}$). As expected if escitalopram level selectively, or at least predominantly, affected state inference, escitalopram level related positively and significantly with P(Go) for the rewarding ($b$ = 0.019, $z$ = 2.97, $p$ = .003, 95% CI [0.006, 0.031], $OR$ = 1.02) and neutral ($b$ = 0.025, $z$ = 3.85, $p$ < .001, 95% CI [0.012, 0.038], $OR$ = 1.03) seasons, but not for the punishing season, in which the relation was negative, albeit not significantly (Supplementary Results Fig. 4; b = –0.006, z = –1.11, p = .268, 95% CI [–0.017, 0.005], $OR$ = 0.99). Moreover, the slopes for escitalopram level differed significantly both when comparing rewarding vs.

punishing seasons (*b* = 0.025, *z* = 4.51, $p < 10^{-5}$, 95% CI [0.014, 0.036], ratio of *OR*s = 1.03) and when comparing neutral vs. punishing seasons (*b* = 0.032, *z* = 5.43, $p < 10^{-7}$, 95% CI [0.020, 0.043], ratio of *OR*s = 1.03). These differential effects of escitalopram level on the different seasons suggest that escitalopram predominantly, if not selectively, affected state inference.

Consistent with the lack of a significant group effect in all analyses in this article, neither group nor the interaction of group by season type were significant ($\chi^2_1$ = 0.62, *p* = .429, and $\chi^2_2$ = 0.44, *p* = .803, respectively).

**Escitalopram did not modulate punishment-based learning or performance**

Serotonin has long been related to punishment[9–11]. In our task, punishments induce passive avoidance, thereby tying together punishment and behavioral inhibition—an intersection that has been especially related to serotonin[9,12–15]. As noted in the main text, however, our model-based analyses did not detect an effect of escitalopram on the parameters that capture punishment-based learning and performance ($\alpha^-$ and $\beta^-$, respectively). We investigated whether we could find evidence that escitalopram (level or group) affected punishment-based learning and performance in our task with two additional analyses that did not use computational models.

In the first analysis, we focused on participants' choices in the punishing seasons that occurred in the first phase (i.e., the first phase of shells PRPR and PNPN; Fig. 1c), because the first phase is not confounded by state inference (Fig. 1d). We used logistic regression, with the counts of Go and NoGo choices in the punishing seasons that occurred in the first phase as the dependent variable, and with group and escitalopram level as independent variables. This regression showed no effect of escitalopram level (*b* = 0.002, *z* = 0.32, *p* = .750, 95% CI [–0.010, 0.014], *OR* = 1.00) or group (*b* = –0.05, *z* = –0.33, *p* = .739, 95% CI [–0.31, 0.22], *OR* = 0.96).

In the second analysis, we tested the idea that serotonin is involved specifically when it is necessary to inhibit a previously rewarded response because it now leads to punishment[16,17]. For that purpose, we tested whether escitalopram (level or group) modulated participants' choices in the second phase of shell RPRP (Fig. 1c), which involves inhibiting a response that was previously rewarded. We used only the second phase because subsequent phases involve state revisiting, so performance in those phases is strongly modulated by state inference. As in the first analysis, we used logistic regression, with the counts of Go and NoGo choices as the dependent variable, and with escitalopram level and group as independent variables. Again, this regression showed no effect of escitalopram level (*b* = –0.002, *z* = –0.21, two-tailed *p* = .830, 95% CI [–0.021, 0.017], *OR* = 1.00) or group (*b* = 0.005, *z* = 0.03, two-tailed *p* = .979, 95% CI [–0.374, 0.384], *OR* = 1.00).

These analyses, like the model-based analyses, found no evidence that escitalopram (level or group) was associated with punishment-based learning or performance. Escitalopram (level or group) did not modulate passive avoidance, even when the punished response had been rewarded previously. Despite the limitations inherent in arguing for the null hypothesis, all the *OR*s in these analyses were very close to 1, which suggests that there really was no effect.

### SUPPLEMENTARY RESULTS REFERENCES

1. Rigoux, L., Stephan, K. E., Friston, K. J. & Daunizeau, J. Bayesian model selection for group studies — revisited. *NeuroImage* **84**, 971–985 (2014).
2. Maia, T. V. & Conceição, V. A. The roles of phasic and tonic dopamine in tic learning and expression. *Biol. Psychiatry* **82**, 401–412 (2017).
3. Daw, N. D. Trial-by-trial data analysis using computational models. *Decision Making, Affect, and Learning: Attention and Performance XXIII* **23**, 3–38 (2011).
4. Wilson, R. C. & Collins, A. G. E. Ten simple rules for the computational modeling of behavioral data. *eLife* **8**, e49547 (2019).
5. Cook, R. D. & Weisberg, S. *Applied Regression Including Computing and Graphics*. (John Wiley & Sons, Inc., 1999).
6. Wiecki, T. V., Poland, J. & Frank, M. J. Model-based cognitive neuroscience approaches to computational psychiatry: clustering and classification. *Clin. Psychol. Sci.* **3**, 378–399 (2015).
7. Portêlo, A., Shiban, Y. & Maia, T. V. Mathematical characterization of changes in fear during exposure therapy. *Biol. Psychiatry Cogn. Neurosci. Neuroimaging* **6**, 1090–1099 (2021).
8. Guitart-Masip, M. *et al.* Differential, but not opponent, effects of l-DOPA and citalopram on action learning with reward and punishment. *Psychopharmacology* **231**, 955–966 (2014).
9. Cools, R., Nakamura, K. & Daw, N. D. Serotonin and dopamine: unifying affective, activational, and decision functions. *Neuropsychopharmacol.* **36**, 98–113 (2011).
10. Daw, N. D., Kakade, S. & Dayan, P. Opponent interactions between serotonin and dopamine. *Neural Netw.* **15**, 603–616 (2002).
11. Dayan, P. & Huys, Q. J. M. Serotonin in affective control. *Annu. Rev. Neurosci.* **32**, 95–126 (2009).
12. Faulkner, P. & Deakin, J. F. W. The role of serotonin in reward, punishment and behavioural inhibition in humans: Insights from studies with acute tryptophan depletion. *Neurosci. Biobehav. Rev.* **46**, 365–378 (2014).
